# Symmetry Breaking under Vectorial Perturbations in Macromolecular X-ray Crystallography

**DOI:** 10.64898/2026.09.28.753211

**Authors:** Harrison K. Wang, Jack B. Greisman, Kevin M. Dalton, Rick A. Hewitt, K. Ian White, Doeke R. Hekstra

## Abstract

Proteins are molecular machines whose dynamics govern their function. Using time-resolved crystallography, the basis of protein mechanics can be uncovered by physical perturbation on protein crystals. Perturbations are often vector fields that, in crystals, break symmetry. Lacking existing tools for analyzing symmetry breaking, we built open-source software to do so. We applied this method to three time-resolved crystallography experiments and analyzed the effect of vectorial perturbations on macromolecular crystal structure. We demonstrate that our method can handle symmetry reduction in all 65 Sohncke space groups.

## I. INTRODUCTION

As molecular machines, proteins undergo series of conformational changes that mediate transport, assembly, and chemical reactions, as well as allosteric modulation thereof. The physical behavior of proteins are complex. Because biochemical stimuli can be described as applying patterns of forces to proteins, it may be possible to simplify our understanding of complex stimuli by observing the response of proteins to controlled patterns of force. Crystals are a unique medium for studying proteins, both defining a regular molecular orientation to physical perturbations and enabling experiments at high temporal and spatial resolution. The natural experiment, then, is to pump a crystal with one of these perturbations, applied as light illumination, temperature, high pressure, or mechanical force, then probe structural changes with X-ray pulses. Time-resolved X-ray crystallography is at the forefront of these pump-probe experiments. For instance, temperature-jump crystallography has uncovered propagation of energy across model enzymes^1^. Additionally, light can activate functional intermediates and interactions in proteins involved in signal transduction^2,3^, oxygen transport^4–6^, ion transport across membranes^7–9^, and DNA repair^10^. Importantly, certain physical perturbations result in symmetry breaking including, for example, a temperature-dependent phase transition in crystals of human dihydrofolate reductase (DHFR)^11^, temperature- and pressure-dependent phase transitions in tetragonal lysozyme^12,13^, and a hydration-dependent phase transition in crystals of Cytomegalovirus immediate-early 1 (IE1) protein^14^. These represent special cases of macromolecular crystallography, where crystal symmetry is broken.

In our studies of protein physics, we previously applied an electric field perturbation to crystals of several proteins using a technique called electric field-stimulated time-resolved crystallography (EF-X)^8,15,16^. These studies revealed patterns of protein conformational change, such as coupling between hinge-bending and active-site remodeling in the *E. coli* dihydrofolate reductase (ecDHFR)^16^; motions resembling ligand binding in PDZ domains^15^; and conductive motions of potassium ions and current-limiting residues in the potassium-selective variant of the bacterial Na-K channel, NaK2K^8^. However, our analysis of diffraction images was often complicated by the fact applying an electric field can change the underlying symmetry of the crystal lattice. That is, different molecules in each unit cell will experience a different relative orientation of the field, leading to differential stimulation (**Figure 1a-b**), thus breaking the crystal symmetry. Indeed, symmetry breaking can happen whenever the physical perturbation is a vector quantity. For example, pressure shockwaves generated by X-ray probes^17^, quantum phenomena resulting from weak terahertz laser pulses^18^, and even forces applied to magnetic dipolar sidechains could possibly constitute vectorial perturbations of the crystal, explained in ref. 19. We note that, although we focus here on symmetry breaking in macromolecular crystals, similar phenomena occur in materials science and chemistry. In our prior work, we encountered various challenges in analyzing symmetry breaking under vectorial perturbations in macromolecular crystallography. We asked whether there was a general formalism for handling vectorial perturbations in such experiments, but to our knowledge, none existed. Still, there is a deep understanding of symmetry breaking due to phase transitions in crystals of small molecules^20–22^. As macromolecular crystals are frequently pseudosymmetric and require analysis in multiple space groups, methods for changing space group basis are also well-developed^23–26^. Furthermore, another form of symmetry breaking is general and well-studied: the case of anomalous scattering, where the Friedel symmetry is broken. Drawing on this body of work, we built a general approach to enable analysis of symmetry breaking due to vectorial perturbations during macromolecular crystallography, and provide corresponding open-source software.

**FIG. 1:**
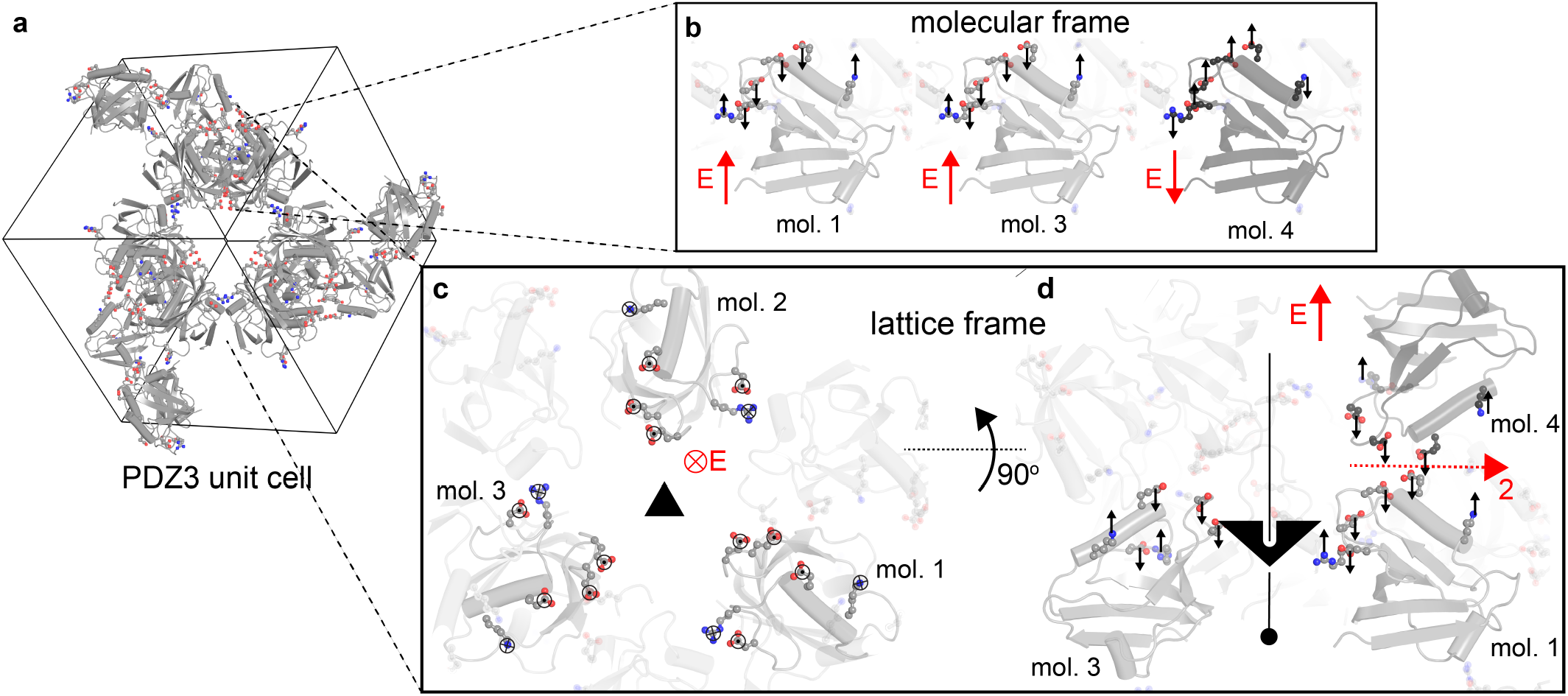
Effect of vectorial perturbations on crystallographic symmetry. An electric field is applied to the crystal lattice of PSD-95 PDZ3^47^. **a)** The crystallographic unit cell contains 24 copies of PDZ3 that differ not by their conformation, but by their orientation. The unit cell is a cube, here viewed along a body diagonal. **b)** The electric field direction in the reference frame of three PDZ3 molecules, labeled mol. 1, mol. 3, and mol. 4. **c)** Three PDZ3 molecules, labeled mol. 1-3, are symmetric by a threefold rotation (black triangle) with an axis pointed into the page. These three copies are also oriented the same to an electric field pointing into the page, so they experience the same pattern of forces (black ⊗ and ⊙) resulting from the electric field (red ⊗). **d)** View rotated 90 degrees from **b**. Molecules 1 and 4 are oriented differently to the electric field, so their electric field response is different (black arrows). A twofold rotation axis, indicated as an arrow labeled with the number 2, previously related molecules 1 and 4 (red dashed line and dyad), but is now broken due to the different electric field responses of molecules 1 and 4. The threefold rotation in **b** is still preserved (black triangle and line). See further handling of PDZ3 symmetry breaking in **Figure 4**.

We applied this method to three examples from time-resolved crystallography. Finally, we show that our method can generalize to all 65 Sohncke space groups.

## II. FORMAL DESCRIPTION OF SYMMETRY REDUCTION

Crystal lattices generally contain multiple copies of a macromolecule in different orientations (**Figure 1a**). When a vectorial perturbation is applied to a crystal lattice, some sets of these copies are oriented the same to the vectorial perturbation, while other sets are oriented differently. Similarly-oriented molecules will respond similarly, while differently-oriented molecules may respond differently (**Figure 1b**). For ease of exposition, we will suppose the vectorial perturbation is an electric field, but the approach is general. How do we systematically determine the orientation of each molecule to the electric field, and each molecule’s subsequent conformational response?

Naively, we could start by inspecting every single molecule in the unit cell and their response to the electric field. However, this is redundant in instances where the unit cell contains many copies of the macromolecule. Instead, we can simplify our analysis by relying on several mathematical observations about crystallographic symmetry. The relative orientation of each molecule in the unit cell is not random but determined by the unique set of symmetry operations corresponding to each space group. When an electric field is applied to a crystal lattice, some symmetries are broken and some are preserved. A symmetry is preserved if it relates two molecules in the unit cell with the exact same response to the electric field (**Figure 1c**). In contrast, a symmetry is broken if it now relates two molecules that do not have the same electric field response (**Figure 1d**). To our main goal of determining the electric field response of each molecule, it would help us to classify molecules by symmetry, and therefore determine the preserved and broken symmetries for a given electric field direction.

To quantify the degree to which a symmetry is broken by a vectorial perturbation pointing along the unit direction 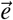, we introduce a metric called the field-symmetry alignment (FSA). This is defined as 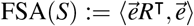, where *R* is the rotation matrix of symmetry operation *S*, and we use the field direction 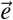 in the fractional basis for ease of rotation, and we also use the 3 by 3 metric tensor *G* = *O*^⊺^*O* (where *O* is the orthogonalization matrix^27^) as the matrix representation of our inner product ⟨·, ·⟩. In essence, instead of the simple inner product 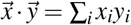inner products in fractional space are 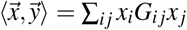. Note that we can ignore the translation component of the symmetry operation *S*, as vectors are invariant under translation. Hereafter, all vectors are treated as row vectors (unless denoted otherwise), and thus transformed by right multiplication. The FSA will aid us in determining the preserved and broken symmetries for a given electric field orientation.

### A. Preserved symmetries

Let *H* be the space group of a macromolecular crystal and *L* be the low-symmetry space group that results after applying a vectorial perturbation to that crystal. As discussed above, it would simplify our analysis to find *L* that contains preserved symmetries. We use two observations to arrive at a choice of *L*. First, we reiterate that *L* must be a subgroup of *H*, as it must both be a group in the algebraic sense^28^, and contain only the symmetries of the parent group^29^ (p. 735). Second, if the field 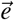 is applied parallel to a symmetry axis, the corresponding symmetry operations are preserved. In other words, 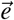 is preserved if and only if the symmetry operation transforms 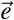 to itself (**Figure 1b**). Three equivalent observations follow: first, for all symmetry operations *L*_*a*_ in *L*, FSA(*L*_*a*_) = 1. Second, 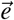 is an eigenvector of each *L*_*a*_. Third, in group-theoretic language, *L* is the stabilizer of 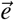. This idea is closely related to the isotropic subgroup in ref. 30, although in our case, the group action is on a vector in ℝ^3^ instead of in the space of all possible atomic displacements. With these two observations, we describe how to choose *L* later.

With this definition of preserved symmetry, we may attempt during the experiment to perfectly align the electric field along a rotation symmetry axis. But this is not simple given the experimental geometry. In general, and in our examples provided below, we do not have perfect alignment. However, if an electric field is misaligned by a small amount, two molecules related by a symmetry operation can still experience essentially the same electric field. Electric field orientations that are off by up to 15° still have FSA of more than 85%, which suggests that preserving a symmetry axis deviating up to 15° from an electric field can still help average together the electric field responses from similarly-responding molecules. Thus, in general and in our examples below, any nontrivial low-symmetry group used in our analyses is an approximation rather than a true symmetry.

### B. Broken symmetries

Any symmetry operation that does not end up in *L* is then a broken symmetry operation. We utilize these broken symmetries to measure the degree of symmetry breaking (which we will describe in detail in Measures of signal) and to compare formerly-equivalent molecules. However, we need not use all broken symmetries at once for analysis, as some of the broken symmetries may be equivalent to each other, i.e., belong to the same coset^28^. Here, notate *R*_*H*_ as the group of non-translational operations of the high-symmetry space group (i.e., removing any unit cell translations or centering operations). Now we make one observation in three equivalent ways. First, if the number of cosets is notated as [*R*_*H*_ : *R*_*L*_], where *R*_*L*_ is a subgroup of *R*_*L*_, then by Lagrange’s theorem:

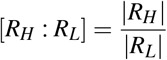

where | · | denotes the order of a group. Second, in group-theoretic language, as in the orbit-stabilizer theorem, [*R*_*H*_ : *R*_*L*_] is the order of the *orbit* of 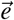, i.e., the number of unique ways that *H* can transform 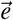. Third, each equivalent symmetry transforms the asymmetric unit equivalently, while each nonequivalent symmetry transforms the asymmetric unit nonequivalently. From these three observations, a set of [*R*_*H*_ : *R*_*L*_] operations from *H* can fully represent the |*R*_*H*_ | − | *R*_*L*_| broken symmetry operations when computing symmetry-breaking maps or statistics, which we will see in **Section IV**. What are the members of each class of equivalent symmetries? It is straightforward to determine the contents of each coset. Denote *H*_*i*_, *H*_*j*_, *L*_*x*_ as rotation group elements (i.e., rotation symmetries) of *R*_*H*_ and *R*_*L*_, respectively. Then *H*_*i*_ is equivalent to *H*_*j*_ if and only if there exists some *L*_*x*_ ∈ *L* such that *H*_*j*_*L*_*x*_ = *H*_*i*_. We show how to partition the eight P422 symmetry operations into cosets in **Figure S1**.

We make three helpful observations about the FSA of broken symmetries. First, if symmetry operation *H*_*b*_ has FSA(*H*_*b*_) = −1, it perfectly inverts the electric field orientation 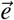. For rotational and screw symmetries, this is possible if and only if there is a 180° rotation around an axis orthogonal to the electric field orientation 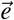. Second, we can see the first observation by first finding a general, closed form for the FSA. If we rotate the field’s unit vector by *ϕ* degrees around some rotation axis, and the unit vector is *θ* degrees away from the axis of rotation, then the FSA is

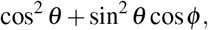

derived in the **Supplementary Information**. This quantity has a unique minimum of −1 at *θ* = *π*/2, *ϕ* = *π*, as stated in the previous observation. Third, elements *H*_*i*_, *H*_*j*_ of the same coset (with rotations denoted also as *H*_*i*_, *H*_*j*_) will have the same FSA. This is because, for some *L*_*i*_ in *L*,

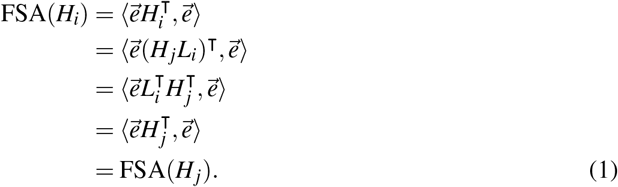

In **Section IV**, the FSA will aid us in identifying the “most broken” symmetry operations.

### C. Basis changes

Electric field perturbation typically preserves the lattice, so molecules are positioned similarly in the low- and high-symmetry space groups. Thus, the unit cell is preserved and can still be used. However, the low-symmetry space group can now be non-canonical due to preserved centering operations (i.e., translational symmetries inside the unit cell) or various space group conventions concerning origin shifts or axis permutations^23–26^. Non-canonical space groups, of which we will see two examples later, accurately describe the lattice but aside from certain cases (such as C1 in GEMMI^31^ or P21 1 1 in CCTBX^32^) are not generally supported by macromolecular crystallography software. In cases where the naive low-symmetry space group is non-canonical, symmetry reduction is required and involves changing the basis of the crystallographic reflections, the unit cell, and the protein model. What follows is a technical discussion that requires some familiarity with crystallographic bases, although we will provide intuition to help. An atomic model of a protein describes atom positions in either a Cartesian or fractional basis. Cartesian positions in the crystal frame are denoted by **X** = (*X,Y, Z*). Cartesian positions are equivalently handled. Atom fractional positions are denoted by **x** = (*x, y, z*) with **X** = **x***A*^*T*^, with *A* the fractional-to-Cartesian basis change matrix to be defined later. Crystallographic reflections are represented by Miller indices, which we notate as **h** = (*h, k, l*) and the inverse-Cartesian positions in reciprocal space determined as **d**\* = **h***A*\*^⊺^. The basis changes discussed here do not affect **X** and **d**\*, as they are objects represented by a Cartesian (or reciprocal-Cartesian) basis in the lab frame, and serve as “solid ground”. Rather than the Cartesian bases, the bases to change are the fractional and reciprocal bases. In essence, we change the unit cell basis vectors **a, b, c** and the reciprocal basis vectors **a**\*, **b**\*, **c**\* into new basis vectors **a**′, **b**′, **c**′, **a**′*, **b** ′*, **c**′*. The original bases can be expressed concisely as 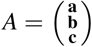 and *A*\*^⊺^ = (**a**\*^⊺^ **b**\*^⊺^ **c**\*^⊺^), respectively the real space and reciprocal space orientation matrices.

We are ready to describe how a basis-change operation *C* affects the crystallographic reflections, the unit cell, and the protein model. We start with changing the basis vectors *A*. We define *C* as the affine operation which transforms *A*^⊺^ to *A*^′⊺^ := [**a**^′⊺^ **b**^′⊺^ **c**^′⊺^] = *C*^⊺^*A*^⊺^. From this, we can already calculate the new unit cell side lengths *a*′,*b*′,*c*′ and angles *α*′,*β* ′, *γ*′ by the standard relations

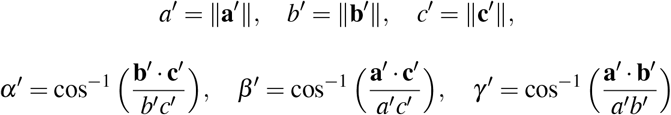

with · as the standard dot product.

Now **X** is conserved regardless of basis choice, so we can use this fact to show how fractional coordinates are transformed:

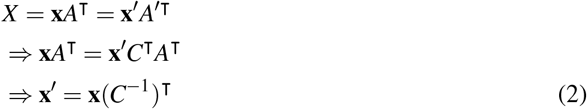

Note that *C* can contain origin shifts, i.e., translations, so it is an affine operation and not a simple 3×3 matrix. Thus, we must handle its inverse carefully when dealing with translations in real space.

So we know how *C* transforms the unit cell and fractional coordinates in real space. What about in reciprocal space? Since *A*^−1^ = *A*\*^⊺^ and *A*′^−1^ = *A*′*^⊺^, we use *A*′ = *AC* = (*A*\*^⊺^)^−1^*C* to find *A*′*^⊺^ = (*A*′^−1^)^⊺^ = *C*^−1^*A*\*^⊺^.

Now, we are ready to transform **h**. Using the fact that **d**\* is conserved regardless of basis choice:

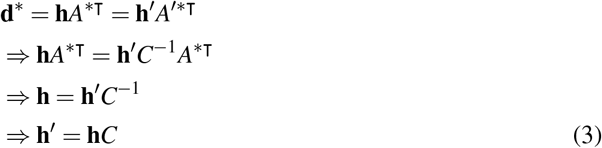

Now we know how the Miller indices **h** of the data are transformed!

How do broken symmetry operations from the old basis work in the new basis? Suppose we have a symmetry operation *S* that transforms **h** to **h**_*S*_ as **h**_*S*_ = **h***S*. We must have **h**′_*S*_ = **h**_*S*_*C*, so **h**′*S* = **h***SC* and **h**′*S* = **h**′*C*^−1^*SC* ⇒ **h**′_*S*_ = **h**′*C*^−1^*SC*. So the operation *S*′ that correctly transforms **h**′ to **h**′_*S*_ is *C*^−1^*SC*. To check, we want to determine whether *S*′ correctly transforms **x**′, i.e., **x**_*S*_ = **x**(*S*^−1^)^⊺^ ⇔ **x**′_*S*_ = **x**′(*S*^′−1^)^⊺^. This is straightforward to verify and we will not show this here. Note that *S* is also an affine operation and not a simple 3×3 rotation matrix.

We are now mathematically equipped to transform high-symmetry maps and models into low-symmetry maps and models. We now draw on computational tools that implement these statements. Basis change operators *C* are unique for a given pair of space groups, derived in ref. 33, well-defined in ref. 29, and implemented in sgtbx. Transformations using basis-change operations are additionally implemented in GEMMI^31^ and reciprocalspaceship^34^. With these computational tools, we are ready to process symmetry-broken data.

## III. PROCESSING SYMMETRY-BROKEN DATA

With the observations of the previous section in hand, we are ready to process symmetry-broken diffraction data. To get from images to a refined model, as with conventional crystallography, we must index, integrate, scale, and merge the data, and then refine the resulting structure factor amplitudes to a low-symmetry model. We modify this workflow to handle symmetry-breaking experiments, as outlined in **Figure 2**. An example of workflow use is found in the regroup Github repository, with supporting information in **Section IV** and the Zenodo deposition associated with this manuscript.

**FIG. 2:**
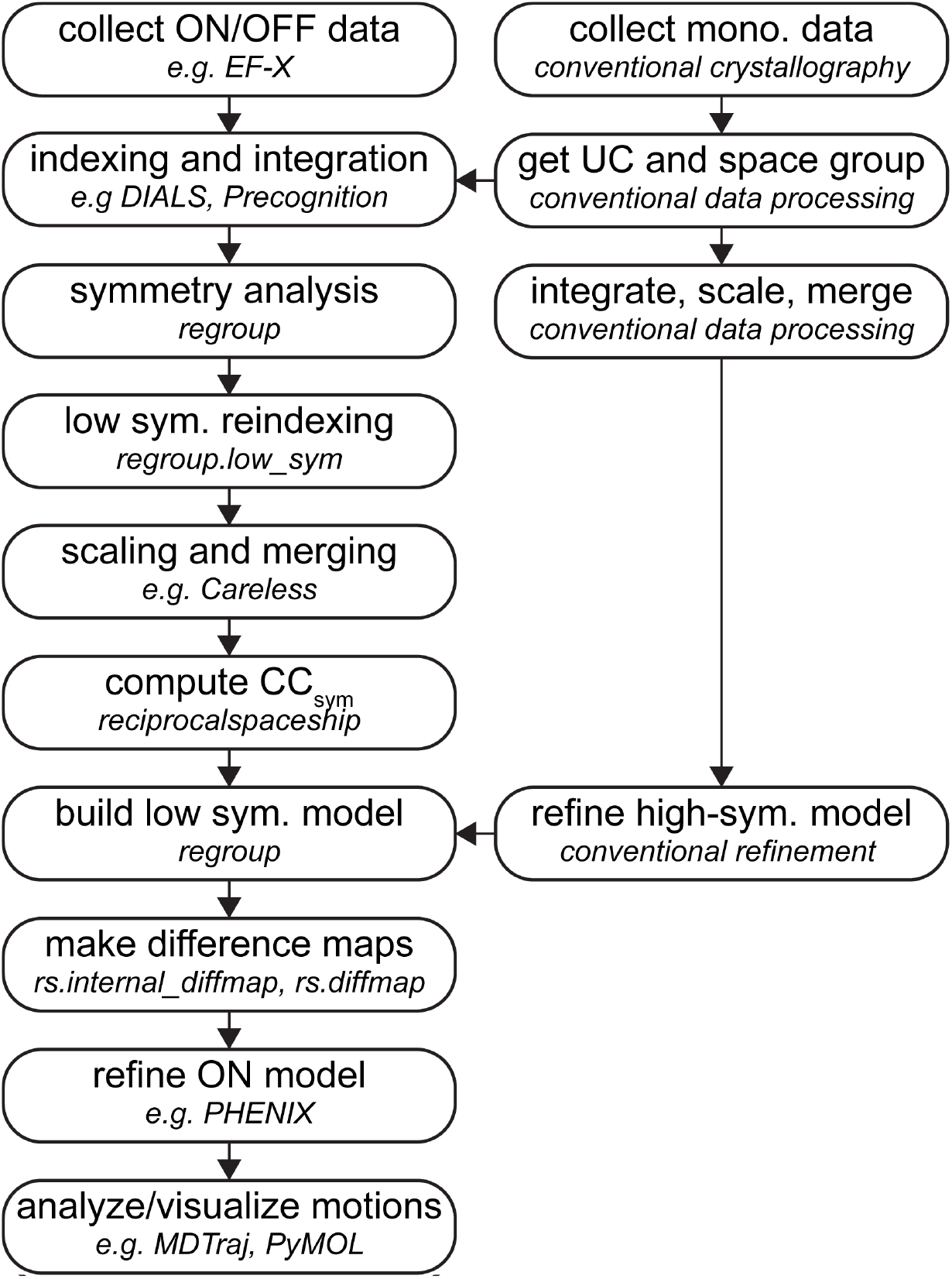
Practical workflow for processing data with vectorial perturbations. Data is collected on a crystal undergoing a vectorial perturbation. These data are indexed and integrated in any format, preferably DIALS, Laue-DIALS^48^, or Precognition. The symmetry is reduced using `regroup`. The data are scaled and merged using Careless. The data are then analyzed with tools from `reciprocalspaceship`. For example, CC_sym_, the degree of symmetry breaking, is measured. If the data are symmetry broken, then a low-symmetry model is constructed using regroup and then refined using Phenix^49^. The coordinates are then be analyzed appropriately. In parallel, a high-resolution dataset is collected on the same crystal construct, and is analyzed conventionally, providing unit cell (UC), space group, and atomic position information.

To begin, we recommend obtaining a high-quality unit cell, space group, and model in the high-symmetry space group by collecting a high-quality monochromatic dataset on crystals at the same data collection temperature and growth condition as those used for symmetry-breaking experiments. Alternatively, the same information can be obtained from the unperturbed control dataset typical to time-resolved experiments. This provides important ground truth for the high-symmetry unit cell, space group, and asymmetric unit (ASU). Upon acquiring symmetry-broken data, we then discuss how to obtain the low-symmetry unit cell, space group, and refined model.

### A. Indexing and integration

As with conventional crystallography, once we have diffraction images, we first index and integrate our symmetry-broken data. Indexing and integration involve assigning observed Miller indices **h**_obs_ := *h*_obs_, *k*_obs_, *l*_obs_ and intensities to each spot in a diffraction image. During data collection, in our experience, spot positions do not typically change, so we can perform indexing and integration in the high-symmetry space group. We are aided by the fact that most indexing and integration programs do not encode any symmetry into the observed Miller indices. However, if the high-symmetry space group contains any screw axes, integration will skip systematic absences due to screw symmetry. Integration in the low-symmetry space group, using any reindexed crystal geometry, will catch these systematic absences. However, we have yet to encounter a case where it is critical to do this (for further discussion of systematic absences, see **Section III G**). After this step, we have the orientation of the crystal lattice in the lab frame as well as a table of unmerged intensities by Miller index.

### B. Inferring the reduced-symmetry space group

During indexing, we also obtain the orientation of the crystal lattice in the lab frame, the *A* matrix, which enables us to determine the direction of the electric field and assign a working low-symmetry space group.

To do this, we developed regroup, a software package built on cctbx^32^ and GEMMI^31^ for identifying suitable low-symmetry subgroups of high-symmetry space groups. As input, regroup takes an orientation matrix *A* inferred from experimental geometry, either from DIALS^35^ or Precognition (Renz Research, Inc.), and the perturbation direction in the lab frame (+*y* in the case of EF-X at BioCARS). `regroup` then identifies the orientation of various symmetry axes in the crystal, and determines the angle between each symmetry axis and the electric field. If the electric field direction is close to any symmetry axis, the low-symmetry space group can be assigned as retaining that symmetry axis. `regroup` also provides a basis-change operation when appropriate. This basis-change operation is sufficient for the next step: reindexing the observed reflections and generating a template low-symmetry model for refinement.

We next describe the details of the regroup algorithm for determining the low-symmetry space group. To begin, we must determine whether each symmetry axis in the crystal is aligned with the electric field direction. If the symmetry axis is parallel to the electric field direction, then the symmetry is aligned and preserved. We can determine the orientation of the symmetry axis by simply determining the real eigenvector of the rotation matrices of each symmetry operation. While this is straightforward to implement, we present another general algorithm that is more intuitive for the EF-X experiment. We start with two observations about crystal geometry. First, crystal facets, a macroscopic observable, can be mathematically related to the unit cell symmetry axes. Symmetry axes are often normal to crystal facets. This detail is especially helpful during the EF-X experiment, in which the electrodes sit flush against a crystal facet. However, regardless of the experiment, this algorithm can be used. Second, the symmetry operations of a suitable subgroup of the high-symmetry spacegroup must all preserve the facet normal orientation (i.e., have FSA=1 with respect to the facet normal). Using these details, we use the following algorithm:

1. Generate the subgroups of the high-symmetry space group using sgtbx^32^. In sgtbx, each subgroup object encodes a basis-change operation where necessary.
2. Generate all possible facets (**h**) up to some absolute index value, and exclude (0,0,0). By default, this × is 1, in which case {(**h**)} is {−1, 0, 1} × {−1, 0, 1} × 1, {−0, 1}, where is the Cartesian product.
3. Compute facet normals 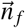 in fractional coordinates as 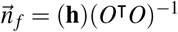 (see ref. 27).
4. For each subgroup:
  - For each subgroup operation, find its rotation matrix *R* (as in ref. 32).
  - Compute whether 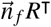 is identical to 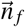.
  - If true for all operations in the subgroup, then this subgroup preserves the facet normal orientation.
5. Find the subgroup with the most preserved symmetry operations. This is the subgroup assigned to (**h**).
6. For each facet, convert the normal vector to the lab (Cartesian) frame as 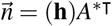 (see ref. 27). The *A* matrix is computed with the lab frame chosen with *z* pointing down the beam, *y* pointing to the floor, and *x* pointing to the center of the synchrotron^36^. As the facet normal corresponds to some symmetry axis, we now have the orientation of each symmetry axis in the lab frame.
7. Find the angle between 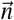 and 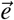 in the lab Cartesian frame, and choose the low-symmetry subgroup with the smallest angle between the facet normal and the electric field orientation.

Now we have the low-symmetry subgroup with one or more preserved symmetry operations. We find that this algorithm works well for the EF-X experiment. Thus far, we have not determined the electric field direction in real space. This would be useful for visualizing protein models, so for a given facet normal, we can convert the facet’s Miller index **h** to the crystal Cartesian frame as **e**_**c**_ = **h***O*^−1^. This computation is implemented as the `efvector.py` pymol script in the `regroup` package.

Once a low-symmetry subgroup is selected, we now know which symmetry operations to consider as preserved, and which to consider broken. We can use these symmetry operations for downstream analysis. As a first example, we can now compute the FSA for each broken symmetry operation. FSA computation is implemented in regroup, enabled with the `--fsa` flag.

### C. Basis change

We now have everything needed to generate low-symmetry unmerged intensities, unit cells, and starting models from their high-symmetry counterparts, up to a basis change. We discuss several instances where a basis change is recommended. We first consider the simple case where the unit cell is unchanged. No reindexing is needed, and we may simply generate the low-symmetry models by using the original symmetry of the crystal lattice.

We outline three more cases where it is not as simple. First, reindexing is sometimes encouraged when reducing the symmetry from a primitive space group to one of its subgroups (e.g., from P2_1_2_1_2_1_ to P2_1_11). This is because of a non-canonical rotation in the unit cell axes, as described in **Section II C**. It is advisable, in this case, to rotate the unit cell axes from the P2_1_11 setting to the conventional *P*12_1_1. Second, reindexing is encouraged when the low-symmetry space group is primitive and the high-symmetry space group is centered, i.e., the space group symmetries include translational operations by fractions of the unit cell. If we use the same unit cell, in the low-symmetry model, we retain copies of the high-symmetry asymmetric unit (ASU) that are related solely by this centering symmetry. However, we only need one copy of each model per field orientation; extra copies add degenerate refinement parameters. In this case, if there are *c* centering operations retained, the reindexed unit cell must be *c* times smaller than the old unit cell, corresponding to *c* times fewer copies of the high-symmetry ASU in the unit cell. Third, reindexing is required when the low-symmetry space group has different centering than the high-symmetry space group, regardless of whether the latter is centered or primitive. This is since centered space groups require a precise choice of origin and setting, which are, in general, not interchangeable. As reindexing can be complicated to understand, we illustrate the latter two cases in **Section IV**.

### D. Low-symmetry maps and models

To obtain the low-symmetry model, we expand the high-symmetry ASU using the broken symmetry operations, and then concatenate each unique copy into a single model (**Section II C**). This can then be validated by comparison to a model from molecular replacement of the high-symmetry ASU into the low-symmetry data. In the low-symmetry model, we expect 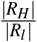 copies of the high-symmetry ASU, where *R* and *R* are defined as in **Section II B**. As for the unmerged intensities, they are denoted by observed Miller indices **h**_obs_ and no further action is required.

In cases where reindexing occurs, the unit cell and unmerged intensities are transformed following **Section II C**. A routine for transforming merged and unmerged intensities is implemented in regroup.low_sym. The low-symmetry model is transformed by a reindexing operation. In all cases, we now have the information to continue processing the data in the low-symmetry space group.

### E. Scaling & Merging

We now seek to compute high- and low-symmetry structure factor amplitudes from scaling and merging the respective unmerged intensities. Naïvely, the structure factor amplitude is the square root of the merged intensity. However, during the crystallography experiment, unmerged intensities may deviate from this relationship by accumulating systematic errors from the experimental geometry, such as from detector aberrations, beam polarization, crystal morphology, fluctuations in the beam intensity, lattice defects, and X-ray absorption by the crystal or surrounding material^37–39^. Both high- and low-symmetry structure factors contain information useful for fitting a scale function that captures these systematic errors, as we know that these reflections were previously identical and will still have similar intensities after symmetry breaking. To maximally extract such information during scaling, we would benefit from simultaneous scaling of multiple ON and OFF datasets in both high- and low-symmetry space groups, while also mapping the relationships between equivalent reflections in the different space groups. To this end, we modified Careless, a scaling routine, to handle relationships between reindexed reflections. Careless is a variational Bayesian inference method that estimates a scale parameter for each intensity without a parametrized physical model^37^. Careless scales reflections as a function of their metadata, such as image number, detector position, and the observed Miller indices **h**_**obs**_. High-symmetry and low-symmetry data can be merged simultaneously in Careless. Careless uses a multivariate Wilson prior^40^ to impose correlation between intensities of the same reflection in two different datasets. We modified Careless to correctly handle correlations between high-symmetry data and their corresponding reindexed, low-symmetry data. To specify this relationship, Careless takes as input a reindexing operation per pair of related datasets. regroup outputs the required reindexing operation by default.

While using Careless for analyzing symmetry breaking, one detail requires special attention. Careless uses reflection metadata to estimate a scale function. We do not recommend naïvely using **h**_**obs**_ as metadata for both the high- and low-symmetry intensities at the same time. This is because while the **h**_**obs**_ of the high- and low-symmetry reflections are related by a reindexing operation, Careless does not automatically reindex these metadata even if the reindexing is specified in the multivariate Wilson prior. Consequently, for any high- and low-symmetry reflections that share the same **h**_**obs**_, Careless handles these reflections similarly even though there is no reason they are related. In certain cases, this leads to poor scaling, as we will see in one example. To repair this issue, we add the high-symmetry **h**_**obs**_ as metadata for the low-symmetry intensities, and then use high-symmetry **h**_**obs**_ instead of the general **h**_**obs**_ as metadata. These metadata can be added using regroup.low_sym. Careless is a powerful but bespoke method, involving training a neural network, controlled by many tunable hyperparameters with optimal values that depend on the dataset. For general tips on using Careless, we refer the reader to references 38–40.

### F. Refinement

We are now ready to determine the conformational response of a macromolecule to each orientation of the electric field. We have structure factor amplitudes that are scaled and merged in the low-symmetry space group, and we also have a low-symmetry starting model. To recover conformational changes, we proceed by simply refining the model against the data using conventional refinement practices in time-resolved crystallography.

### G. Treatment of systematic absences

In reciprocal space, intensities appear or disappear at certain positions in the reciprocal lattice depending on the crystal symmetry, and must be handled with care during indexing. Reflections that disappear are called systematic absences. There are two relevant categories of systematic absences for chiral macromolecules: those arising from centering operations, and those arising from screw axes. We call these centering and screw systematic absences, respectively. We do not expect the electric field to affect the centering systematic absences. This is because two copies of the asymmetric unit related by a centering operation are oriented the same relative to the vectorial perturbation, and the centering symmetry therefore does not break. As a consequence, when we break symmetry, we never expect the reappearance of centering systematic absences. Rather, basis changes will account for centering systematic absences, as we will see in our first example (**Section IV A**).

In contrast to centering symmetries, screw symmetries may break due to electric field. Thus, we do indeed expect the electric field to change whether reflections are screw systematic absences. As a consequence, in principle screw systematic absences should be treated properly during scaling and merging. As of version 0.3.1, when using a multivariate Wilson prior^40^, Careless handles the reappearance of screw systematic absences. We elaborate on how this is handled mathematically. With a multivariate Wilson prior, Careless provides the distribution of each structure factor amplitude of a child dataset, conditioned on each structure factor amplitude in a corresponding parent dataset. Suppose a child reflection *h*_*c*_ with normalized structure factor amplitude *E*_*c*_ depends on a parent reflection *h*_*p*_ with normalized structure factor amplitude *E*_*p*_. Then under the double-Wilson model, 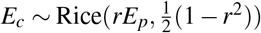. However, when *h*_*p*_ is systematically absent, instead of omitting *h*_*c*_ (i.e., setting *E*_*c*_ = 0), the double-Wilson model has 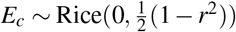^40^.

However, two observations complicate handling screw systematic absences. First, screw operations affect a much smaller portion of reflections, numbering in the tens to hundreds of reflections, or on the order of <1% (typically only *h*00, 0*k*0, 00*l*). Second, in the case of polychromatic data, screw systematic absences are frequently overlapped on top of non-absent harmonics, since the reciprocal lattice points *h*00, 0*k*0, 00*l* each lie on one central ray. This complicates analysis, as measuring the intensity of reappearing screw systematic absences requires harmonic deconvolution. In our experience collecting polychromatic symmetry-breaking data, we have not found an instance where screw systematic absences have meaningfully affected data quality.

### H. Measures of signal

It can be time-consuming to refine a model. Can we tell whether there is meaningful symmetry breaking in the data before refinement? We find it useful to measure the extent of symmetry breaking first in reciprocal space, then in the electron density.

Our most important measure of symmetry breaking borrows a statistic from anomalous diffraction. Unrelated to vectorial perturbations, anomalous scattering leads to Friedel symmetry breaking (while not breaking any space-group symmetry). To evaluate the reproducibility of symmetry breaking of the Friedel operator 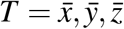, it is common to compute CC_anom_^41^:

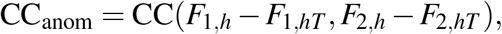

where CC is the correlation coefficient; *F*_*hT*_ is the Friedel mate of *F*_*h*_; and the set of structure factor amplitudes *F* has been split, prior to merging, into random half-datasets with indices 1 and 2. The correlation is calculated over all observed *h* in the reciprocal asymmetric unit. Larger CC_anom_ indicates greater symmetry breaking through anomalous scattering.

By analogy, we introduce CC_sym_, the correlation coefficient of symmetry breaking, which measures the reproducibility of a general symmetry breaking. To illustrate, first consider a high-symmetry dataset *F* in space group *H* with some symmetry operation *S*. We compare the merged structure factor amplitudes *F* between regions of reciprocal space that are related by crystallographic symmetry–denote this *F*_*h*_ and *F*_*hS*_. In the OFF data, even when merging in a low-symmetry space group, this symmetry should be intact, so we expect *F*_*h*_ = *F*_*hS*_ for all *h*. Splitting the data *F* into half-datasets *F*_1_, *F*_2_, we expect *F*_1,*h*_ − *F*_1,*hS*_ and *F*_2,*h*_ − *F*_2,*hS*_ to be zero-centered Gaussian noise^40^. Thus, we expect the half-dataset correlation coefficient between each difference

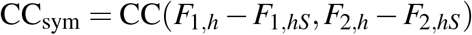

to be 0. Now consider CC_sym_ in the presence of an electric field that breaks symmetry *S*. Since *S* is broken, *F*_*h*_ ≠ *F*_*hS*_ in general. This means datasets collected in the presence of a vectorial perturbation will have nonzero, reproducible *F*_*h*_ − *F*_*hS*_ and yield a positive CC_sym_. We finally note that CC_sym_ calculation requires careful scaling of the data, as spurious CC_sym_ can arise from residual systematic errors (if unaccounted for) that dominate structure factor differences, raising the CC_sym_ baseline.

Second, we rely on canonical real-space methods for estimating signal, such as interpreting ordinary isomorphous difference map peaks. Ordinary difference maps can be affected by crystal heating and radiation damage. To amplify symmetry-breaking signal, we can also generate internal difference maps (*F*_*h*_ − *F*_*hS*_)*e*^*iϕ*^, where *F*_*h*_ and *F*_*hS*_ are again related by a broken symmetry operation *S* ^15^. The real-space equivalent of this operation is subtraction of the electron density of symmetry-equivalent copies of molecules related by *S*. As for phases *ϕ*, we use those from the OFF model in the low-symmetry space group, as obtained in (**Section III D**). In addition, we can threshold weighted isomorphous difference maps at the noise level and can quantify the number of peaks compared to a control isomorphous difference map. Peaks above the noise level in the isomorphous difference maps, consistent with symmetry breaking, are likely to contain signal.

## IV. EXAMPLES

To concretely illustrate concepts from the previous two sections and guide the reader toward a self-consistent understanding of symmetry breaking, we provide three examples where an electric field results in symmetry breaking requiring a basis change. We also draw on our workflow to extract symmetry-breaking signal in crystallographic data.

### A. PDZ2

We first review how to handle the symmetry of the original EF-X experiment^15^ on the second PDZ domain of LNX2 (hereafter PDZ2), a case where we perform a centered-to-primitive basis change (**Section III C**). The unit cell of the PDZ2 crystal is in space group C121, which contains four copies of the asymmetric unit (ASU), a single molecule of PDZ2 (**Figure 3a-b**). In the original experiment, a crystal of PDZ2 was glued to an electrode so that the orientation of the electric field to the crystal lattice could be kept the same throughout the experiment. The electric field was applied along the (1,0,0) facet normal, leading to two distinct electric field responses and thus two molecules of PDZ2 in the low-symmetry ASU (**Figure 3c-d**), while four molecules of PDZ2 in the “C1” unit cell remain. C1 is a noncanonical space group with C-centering and no other symmetry operations. To aid refinement, we then changed the basis to P1 (**Figure 3e-f**), retaining one copy of the low-symmetry in the unit cell. The arrangement of chains in the lattice should be invariant to basis change from C1 to P1. We verify that this is the case (**Figure 3g-i**).

**FIG. 3:**
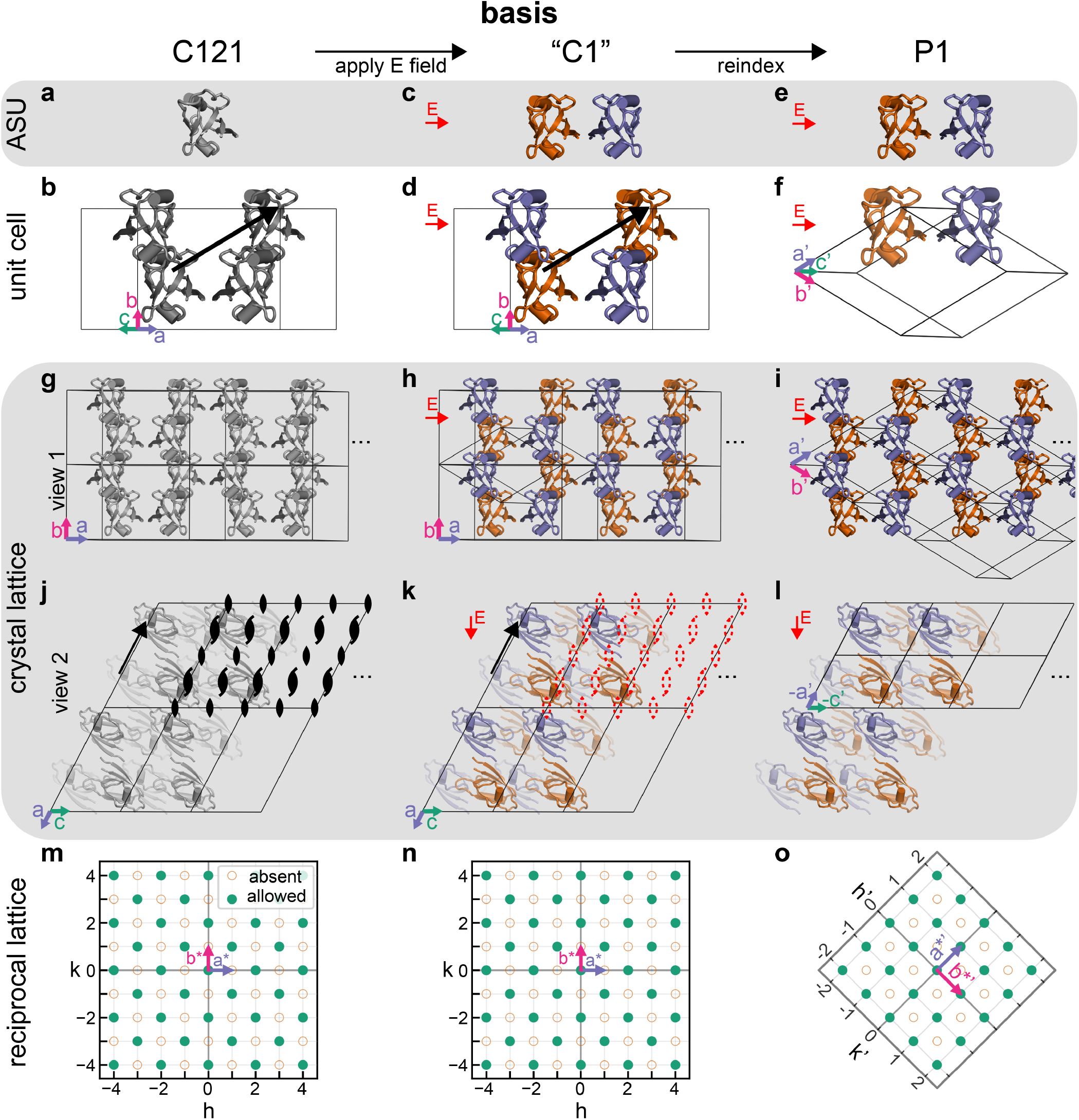
Overview of the basis change during PDZ2 EF-X. **a)** The ASU of LNX2^PDZ2^, hereafter PDZ2, as in ref. 15. **b)** The C121 unit cell of PDZ2, showing four equivalent copies of the asymmetric unit. Copies of PDZ2 are related by centering (black arrow). **c)** The asymmetric unit of PDZ2 in the electric field-excited state, containing two molecules of PDZ2, after applying an electric field normal to the (**h**) = (− 1, 0, 0) facet, as in ref. 15. The space group is now the noncanonical space group “C1”, where centering is preserved. **d)** The unit cell of PDZ2 in C1. There are two asymmetric units of excited-state PDZ2 in the unit cell. Copies of PDZ2 are related by centering (black arrow). **e)** the ASU of PDZ2 after a change of basis. **f)** The new unit cell after a change of basis, containing one copy of the ASU of PDZ2 in the electric field-excited state. In all panels, the electric field vector is shown as a red arrow. Continued on next page. **g-i)** Crystal lattice of PDZ2 in the **g)** C121, **h)** C1, and **i)** P1 space groups. Copies of PDZ2 are related by centering (black arrow). **j-l)** View 2 of the PDZ2 crystal lattice. Symmetry axes in **j)** are shown as black dyads for twofold rotation axes and screw dyads for twofold screw axes. Broken symmetry operations in **k)** are shown as red dashed lines. **m-o)** Plot of the reciprocal grid along *h* and *k* in **m)** C121, **n)** C1, and in **o)** P1, showing that C-centering systematic absences now lie on fractional lattice points in P1. Absent reflections are shown as circles with orange dashed borders. Allowed reflections are shown as green disks. In all panels, the electric field vector is shown as a red arrow.

Why do we do any of this? If we view the ground-state crystal lattice along the *b* axis, we can clearly see the symmetries that relate PDZ2 copies in the C121 unit cell (**Figure 3j**). Twofold rotation and screw symmetries are present throughout the lattice and are parallel to the *b* axis. Centering operations relate copies of PDZ2 in and out of the plane. Due to the rotation symmetries, PDZ2 is in two distinct orientations to the electric field. Thus, the electric field breaks these rotation symmetries, while preserving the centering symmetries (**Figure 3k**). Because of centering, there are two molecules per electric field orientation in the unit cell, and two copies of the ASU per unit cell. As discussed in **Section III C**, this is redundant; it is better to refine one molecule per electric field orientation (unless centering symmetry is somehow enforced during refinement). Thus, we remove the centering operation by reindexing the lattice following the (*C*^−1^)^⊺^ = *a* + *b, a* − *b*, −*c* basis change provided by regroup. It is most illustrative to first change the basis in reciprocal space, as derived in **Section II C**: *A*′*^⊺^ = (*C*^−1^)^⊺^*A*\*^⊺^ so **a**′* = **a**\* + **b**\*, **b**′* = **a**\* − **b**\*, **c**′* = −**c**\*, while **h**′ = **h***C* so *h*′ = (*h* + *k*)/2 and *k*′ = (*h* − *k*)/2. We can directly verify these relations using diagrams using reciprocal space diagrams in the *hk* plane (**Figure 3m-o**), and see that the basis change does not, by itself, affect the spacing between each lattice point. We can also verify the basis change in real space. The unit cell basis vectors **a, b, c** transform to **a**′ = (**a** + **b**)/2, **b**′ = (**a** − **b**)/2, **c**′ = −**c**. In this case, `regroup` provides a basis change that preserves the lattice and reciprocal grid arrangement while changing the reciprocal and real space basis vectors.

Moreover, we see that systematic absences have been properly handled. In C121, there are both screw and centering systematic absences. The centering systematic absences are at (*h, k, l*) | *h* + *k* = 2*n* + 1, *n* ∈ ℤ (**Figure 3m**). Applying an electric field does not lead to reappearance of these reflections. After reindexing, the systematic absences now fall on fractional *h, k* lattice points (**Figure 3o**). The screw systematic absences in C121 are at {(0, *k*, 0) | *k* = 2*n* + 1, *n* ∈ Z}, and the screw symmetry breaks when reducing symmetry to P1. However, in this case, the screw systematic absences are also centering systematic absences, which are still preserved after symmetry breaking, so they do not reappear. To provide a contrasting example, in the space group P2_1_, the screw systematic absences indeed reappear when reducing symmetry to P1 (**Figure S4**).

### B. PDZ3

We next present two new EF-X experiments that illustrate how a vectorial perturbation breaks crystallographic symmetry. The second EF-X experiment consisted of one crystal of the third PDZ domain of PSD-95 (PDZ3), a case where we perform a primitive-to-centered basis change (**Section III C**). Unit cells of PDZ3 belong to space group P4_1_32 and contain 24 copies of the asymmetric unit (**Figure 4a**), each consisting of one copy of PDZ3 (**Figure 4b**). One PDZ3 crystal was mounted on an electrode and high-voltage (ON) and no-voltage (OFF) data were collected on it. To apply the E field in a consistent orientation relative to the crystal lattice during the course of the experiment, we glued the PDZ3 crystal to one electrode while allowing free rotation of this electrode and crystal around an axis parallel to the electric field, while fixing the position of another electrode and the remaining experimental hardware. To interpret the data from this experiment, we followed the workflow outlined in **Figure 2**. To obtain the ground-truth unit cell dimensions, space group, and atomic model of PDZ3 in the same crystal form, we collected a room-temperature dataset of PDZ3 to 1.35 Å, finding a cubic unit cell in space group P4_1_32. With this unit cell in hand, we indexed and integrated these data using Precognition (Renz Research, Inc.). Using regroup, we found that an electric field was applied close to the (1,-1,-1) facet normal, causing all symmetries to break, except for a threefold rotation axis (**Figure 4c**). The low-symmetry space group was assigned to R3:H. Since this space group is R-centered while the original space group was not centered, we needed to change the crystallographic basis.

**FIG. 4:**
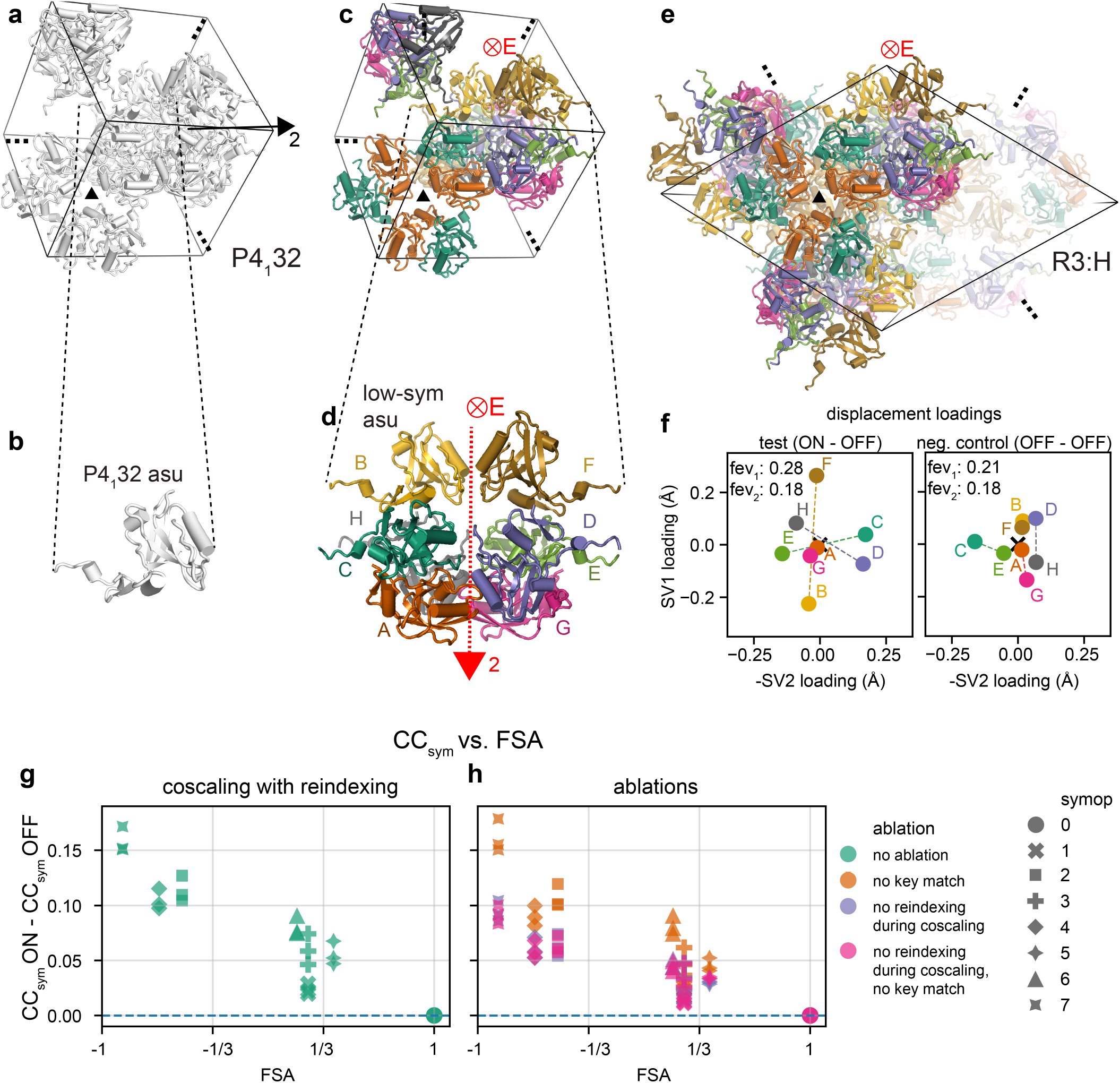
The effect of the electric field direction on PDZ3 motions. **a)** The P4_1_32 unit cell of the third PDZ domain of PSD-95, hereafter PDZ3. A threefold rotation symmetry axis is indicated by a black triangle, and a twofold rotation symmetry axis is indicated by a black dyad and line segment. **b)** The P4_1_32 asymmetric unit (asu) of PDZ3, containing one copy of PDZ3. **c)** The unit cell of PDZ3 after applying an electric field normal to the (**h**) = (1, −1, −1) facet (red ⊗, i.e., into the page). The electric field is parallel to a threefold rotation symmetry axis. **d)** The asymmetric unit (ASU) of PDZ3 in the electric field-excited state, containing eight molecules of PDZ3, each oriented differently to the electric field. Dashed red line and dyad indicate a twofold rotation axis that is broken due to the electric field. Molecules A and G, B and F, C and E, and D and H are oppositely-oriented to the electric field, as in **d). e)** PDZ3 unit cell after a change of basis from P4_1_32 to R3:H. Continued on next page. **f)** Principal component analysis of PDZ3 electric field-induced motions. We performed singular value decomposition on the *Cα* − *Cα* distances across the eight PDZ3 molecules in the low-symmetry ASU, as in ref. 16. Singular vectors (SVs) 1 and 2 per-molecule loadings, for refinement against ON-OFF extrapolated structure factors as well as against OFF-OFF extrapolated structure factors. The electric field induces motions in opposite directions for oppositely-oriented molecules. **g-h)** Difference in the ON and OFF correlation coefficients of symmetry breaking (CC_sym_) for the eight cosets of symmetry operations, numbered 0-7. A key for these symmetry operations can be found in **Figure S2**. The CC_sym_ difference is plotted against the field-symmetry alignment (FSA) for each of the symmetry operation classes. CC_sym_s are measured on data merged with **g)** typical Careless settings as well as data merged with **h)** Careless settings ablated as indicated (**Methods**). “Normal” indicates that scaling with Careless was correctly performed as described in **Section II**. “No key match” indicates that during scaling, we use **h**_**obs**_ metadata, which are not guaranteed to match between corresponding high- and low-symmetry reflections, rather than the correct high-symmetry **h**_**obs**_, which are guaranteed to match between corresponding high- and low-symmetry reflections, as in **Section III E**. “No reindexing ops” indicates that there is no prior correlation imposed between corresponding high and low-symmetry reflections during scaling, as in **Section III E**.

Using the basis-change operation from regroup, we reindexed the observed Miller indices prior to scaling, and we also generated a low-symmetry model. The unit cell changed from *a* = *b* = *c* ≈ 90.5 Å, *α* = *β* = *γ* = 90°, to *a* = *b* ≈ 128 Å, *c* ≈ 157 Å, *α* = *β* = 90°, *γ* = 120°, consistent with a hexagonal unit cell setting. This unit cell has a threefold higher volume, which corresponds with an R-centered space group having three times as many centering operations (including the trivial centering operation). To generate a low-symmetry model, we placed 8 copies of the high-symmetry ASU into the low-symmetry data by symmetry expansion using PyMOL, validated against molecular replacement (**Figure 4d**). To check this result, we verified against our formalism. As three rotation symmetries were retained (including the trivial rotation), by Lagrange’s theorem, PDZ3 experiences 24/3=8 different field directions in the crystal, with the asymmetric unit containing eight formerly-equivalent copies of PDZ3. To ensure that our low-symmetry data is consistent with the crystal symmetry, we tabulate all copies of PDZ3. If we expect 8 copies from the broken symmetries, times 3 copies from the preserved symmetries, times 3 copies from centering, then we expect a total of 8 ∗ 3 ∗ 3 = 72 copies of the high-symmetry asu in the R3:H unit cell. We see that this is the case (**Figure 4e**), and we also see that the lattice arrangement of PDZ3 molecules has not been affected. Following our checks for the low-symmetry map and model, we next scaled and merged the data in Careless with a multivariate Wilson prior encoding two relationships: one between the OFF data in P4_1_32 and the OFF data in R3:H, and one between the OFF data in P4_1_32 and the ON data in R3:H. We measured nonzero CC_sym_s in the ON data and near-zero CC_sym_s in the OFF data, indicating that there was significant symmetry breaking with minimal systematic errors (**Figure S2**). To visualize the effects of symmetry breaking, we then used the most broken symmetry operation, with a FSA of −1, to generate an internal difference map (**Section S6**, **Figure S3**). We found, in this case, that internal isomorphous difference maps were much better at detecting field-dependent structural changes than ordinary isomorphous difference maps (**Figure S3**).

The data were of high-enough quality that we could observe rotamer flips. For certain orientations of the electric field, we found a shift in the rotamer state in several sidechains, revealing perturbation in a network of hydrogen bonds. However, in other orientations of the electric field, it was possible to shift the rotamer state for several subsets of residues, indicating that this hydrogen bond network could decompose into smaller components depending on field direction.

We next sought to measure the response of the PDZ3 backbone to the electric field by refining the low-symmetry model against the electric field-ON data (see **Methods**). This produced an electric field-ON model. To gain further insight into the electric field-dependent motions, we measured the degree to which molecules oriented oppositely to the electric field would also respond oppositely. We note that low-symmetry ASU molecules A and G, B and F, C and E, and D and H respectively experience the electric field in opposite directions (as each pair was formerly related by a twofold rotation orthogonal to the electric field direction, **Figure 4d**). To measure the conformational response to the field, we performed SVD on the field-induced pairwise C*α*-C*α* distances across all eight chains. We found that oppositely-oriented molecules had similar but opposite loadings of the two top-ranked singular vectors. To examine whether this could possibly be due to the data processing alone (perhaps due to SVD artifacts or refinement target dependence on crystal position), we repeated the same SVD analysis on a model refined against (OFF, low-symmetry) minus (OFF, high-symmetry) data. In this model, we see no reversal of motions due to field reversal, as in the ON-refined model. This shows our ON-refined model has captured true electric field-dependent motions. Thus, PDZ3 molecules oppositely-oriented to the electric field responded oppositely (**Figure 4f**).

We might expect the greatest differences in electric field-dependent motions in two molecules that experience the opposite electric field to one another. An equivalent statement in reciprocal space is that we might expect the highest signal-to-noise ratio in the differences between *F*_*h*_ and *F*_*hS*_ for a symmetry operation that maximally inverts the electric field direction. To test this, we measured the correlation between a given symmetry operation’s FSA and the CC_sym_, finding this value was less than zero. Though there is no mathematical equivalence between the FSA and the CC_sym_, a correlation is still expected, as a symmetry operation *S* with a more negative FSA would reorient the electric field orientation more relative to the molecules formerly related by the symmetry operation (as in **Figure 1**). Such a symmetry operation would result in greater electric-field differences in the corresponding symmetry-related structure factor amplitudes, resulting in stronger negative correlations. Accordingly, we found that in the case of PDZ3, the more *S* transformed the electric field, the more robust the *F*_*h*_ − *F*_*hS*_ differences (**Figure 4g**).

Careful scaling is critical for uncovering symmetry-breaking effects. Here, we use Careless for scaling, which models the scale factor as a slowly-varying function of the observed Miller indices. If scaling is not meticulous, symmetry breaking effects can be falsely positive, resulting from residual systematic errors that vary with the crystal symmetry, or falsely negative, resulting from removal of true symmetry breaking through scaling. We tested the effect of various scaling hyperparameters (as described in **Section III E**) on the systematic errors and degree of symmetry breaking, by measuring the effect of their ablations on CC_sym_s. Removing the prior correlation between the reindexed low-symmetry and high-symmetry reflections substantially degraded CC_sym_s (“no reindexing during coscaling”), although if we additionally handled the Miller index metadata naively, we did not further degrade the CC_sym_s (“no key match”, **Figure 4h**). This indicates that for the PDZ3 dataset, the Miller index metadata does not seem to influence scaling compared to the prior correlation, while the prior correlation matters substantially. We will see another instance below where the opposite is the case. Finally, if we naively handled space group conversion without proper reindexing, the CC_sym_s degrade.

### C. HRas

Our third EF-X example involves another centered-to-primitive basis change. HRas, used for EF-X, crystallized and indexed in the R32:H space group (**Figure 5a**). Each asymmetric unit in the unperturbed state contained one copy of HRas (**Figure 5b**). We performed two separate EF-X experiments on HRas. In both experiments, HRas crystals were mounted on electrodes such that when an electric field was applied close to the (−1,0,−1) facet normal, all six rotational symmetry elements were broken (**Figure 5c**). The low-symmetry space group is naively assigned to the noncanonical “R1:H”, which still maintains the centering and setting of R32:H but not any of the rotational symmetries. To convert from R1:H to P1, we applied the basis-change operation specified by regroup, which changed the unit cell from the hexagonal *a* = *b* ≈ 90 Å, *c* ≈ 136 Å, *α* = *β* = 90°, *γ* = 120°, to the rhombohedral primitive *a* = *b* = *c* ≈ 69 Å, *α* = *β* = *γ* ≈ 81.5°. This is accompanied by a three-fold reduction in the unit cell volume, for the same reasons as in the PDZ2 example. Consistent with the five nontrivial broken symmetry operations plus identity, the low-symmetry ASU contained 6 copies of the high-symmetry ASU (**Figure 5c-d**). We scaled and merged data to 1.9 Å and measured CC_sym_ (**Figure 5e**). As before, in both HRas datasets the overall CC_sym_ was negatively correlated with the FSA, except for the symmetry operation with the least FSA. This is unexpected and suggests HRas is much more sensitive to fields in one direction than another, although the quality of either dataset was insufficient to verify this through refinement of an excited-state model (**Tables S4 and S5**).

**FIG. 5:**
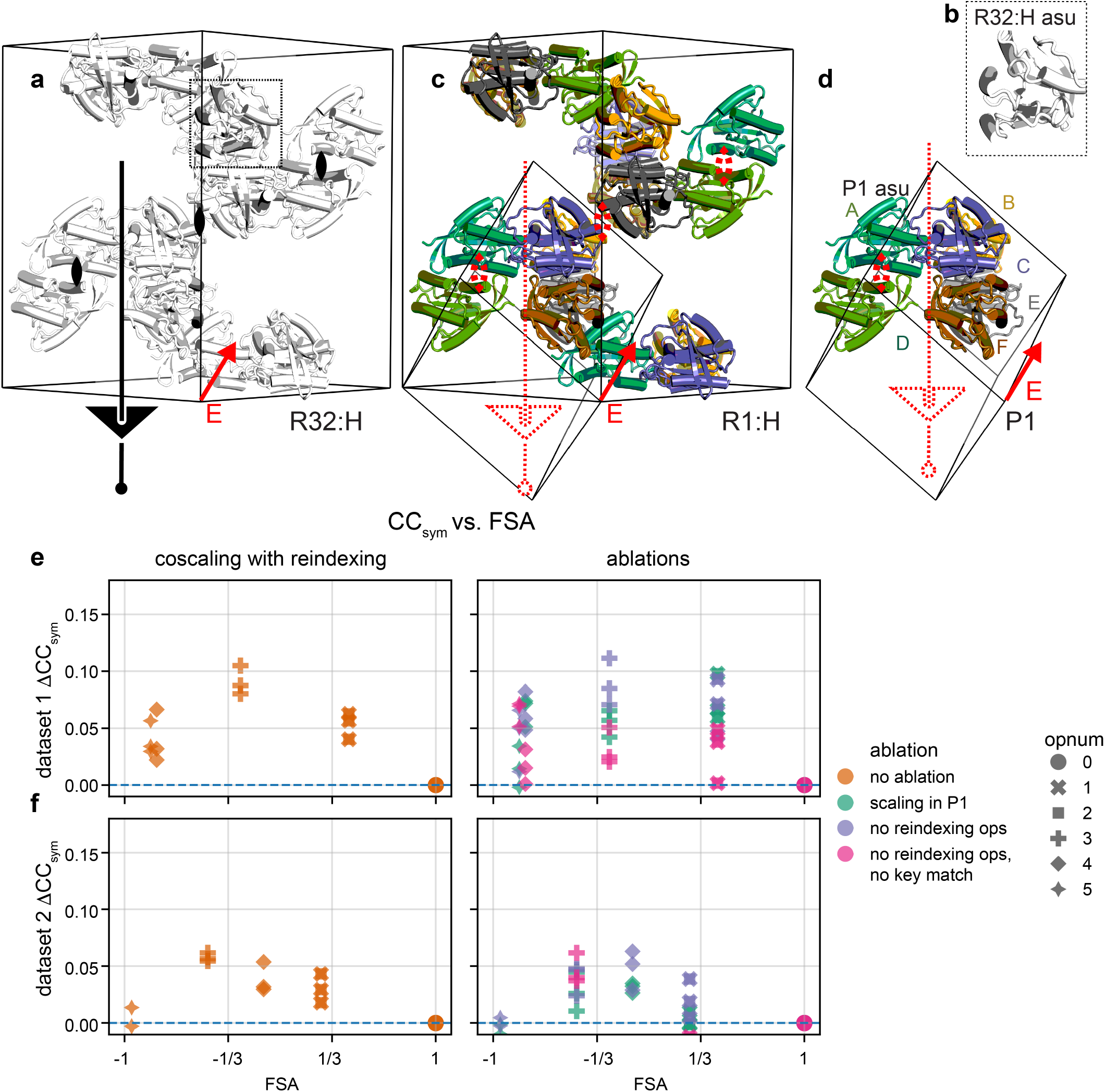
The effect of the electric field direction on HRas motions. **a)** The R32:H crystal lattice of HRas. Black line and triangle indicate a threefold rotation axis. Black dyads indicate twofold rotation axes, viewed on-axis. **b)** Inset of **a)**. The R32 asymmetric unit (ASU) of HRas, containing one copy of HRas. **c)** The noncanonical unit cell of HRas after applying an electric field normal to the (**h**) = (−1, 0, −1) facet (red arrow). **d)** The ASU and P1 unit cell of HRas in the putative electric field-excited state, containing six molecules of HRas, each oriented differently to the electric field. Dashed red line and triangle indicate a threefold rotation axis that is broken due to differential responses of HRas the electric field as a function of orientation. Dashed red dyad indicates a twofold rotation axis that is broken due to differential responses of HRas to the electric field. The electric field direction is indicated in red. Continued on next page. **e)** Δ CC_sym_, the difference in the ON and OFF (CC_sym_) for the six cosets of symmetry operations, numbered 0-5. A key for these symmetry operations can be found in **Figure S6**. This quantity is plotted against the field-symmetry alignment (FSA) for each of the symmetry operation classes. CC_sym_s are measured on data merged with normal Careless settings as well as data merged with Careless settings ablated as indicated. “Scaling in R1:H” indicates that, incorrectly, no change of basis was applied prior to scaling with Careless. “Scaling in P1” indicates that scaling with Careless was correctly performed as described in section **Section II**. “No reindexing ops” indicates that there is no prior correlation imposed between corresponding high and low-symmetry reflections during scaling, as in section **Section III E**. “No key match” indicates that during scaling, we use **h**_**obs**_ metadata, which are not guaranteed to match between corresponding high- and low-symmetry reflections, rather than the correct Hh, which are guaranteed to match between corresponding high- and low-symmetry reflections, as in section **Section III E**.

To test the effect of scaling on CC_sym_s, we modified hyperparameters from standard procedure (**Sections III E and IV B**). First, removing the prior correlation between the reindexed low-symmetry and high-symmetry reflections did not seem to matter (“no reindexing ops”). Second, naively handling the Miller index metadata led to degraded CC_sym_s (“no key match”, **Figure 5e**). Finally, naively handling the space group conversion without reindexing properly (“scaling in R1:H”) also led to degraded CC_sym_s. We observed the same phenomena in a second HRas EF-X dataset (**Figure 5f**). In that dataset, we found that naively handling the Miller index metadata resulted, in fact, in higher OFF CC_sym_s compared to the ON CC_sym_s, which is indicative of poor scaling (**Figure S6**). Taken together, for the HRas datasets, the prior correlation does not seem to matter much for scaling compared to the Miller index metadata, even though the opposite was true for the PDZ3 datasets. These observations illustrate the importance of handling symmetry correctly when scaling.

### D. General case

To test whether we could accurately account for vectorial perturbation applied in a general direction to a general Sohncke space group, we wrote an automated routine for low-symmetry model generation, and checked it against simulated symmetry breaking data from the Protein Data Bank (PDB, see **Methods**). We also tested regroup.low_sym for low-symmetry map generation. Note that this simulation does not add any symmetry-breaking signal, merely just checks that low-symmetry maps and models are accurately generated from corresponding high-symmetry data. Our test set was a random set of 200 protein, DNA, and peptide entries in the PDB across all Sohncke space groups and a small campaign of non-Sohncke space groups. To each dataset, we applied a simulated vectorial perturbation normal to the (1,0,0), (0,1,0), (1,1,0), and (1,-1,-1) facets, and determined the low-symmetry space group, model, and dataset. As a first check, we used this method to recover the HRas and PDZ3 low-symmetry ASUs that we earlier built manually. To validate the low-symmetry models, we rigid-body refined them against the low-symmetry data and measured R-factors. After filtering out 10 PDB entries that did not refine correctly even in the high-symmetry case, we sought to identify successful symmetry reduction. This was defined by our expectations that successful symmetry reduction would neither substantially increase the R-factor or decrease the completeness. We succeeded in all but one case (**Figure 6**). In the one failure, the low-symmetry R factors increased by around 0.06, and high-symmetry data were in the centrosymmetric space group C12/c1. To inspect the cause of the failure, we inspected the low-symmetry model. In each case, the low-symmetry case superposed perfectly on the high-symmetry unit cell. Since the model was generated correctly, the issue lies within the data or the refinement software. We also note that in two of the 190 successes, the high-symmetry data were already below 50% complete^42,43^. We find that the method is robust in all other cases. Thus, we conclude it is possible to reduce the symmetry of macromolecular crystallography data. To extend this work and comprehensively account for all non-Sohncke spacegroups, we note that implementations of subgroup transformations can be borrowed from existing software for handling small-molecule symmetry^26^. We finally note that we did not manually inspect most of the models and there may still be small errors in the generated low-symmetry models. It is straightforward and good practice to validate the generated model by molecular replacement of copies of the high-symmetry ASU into the low-symmetry data.

**FIG. 6:**
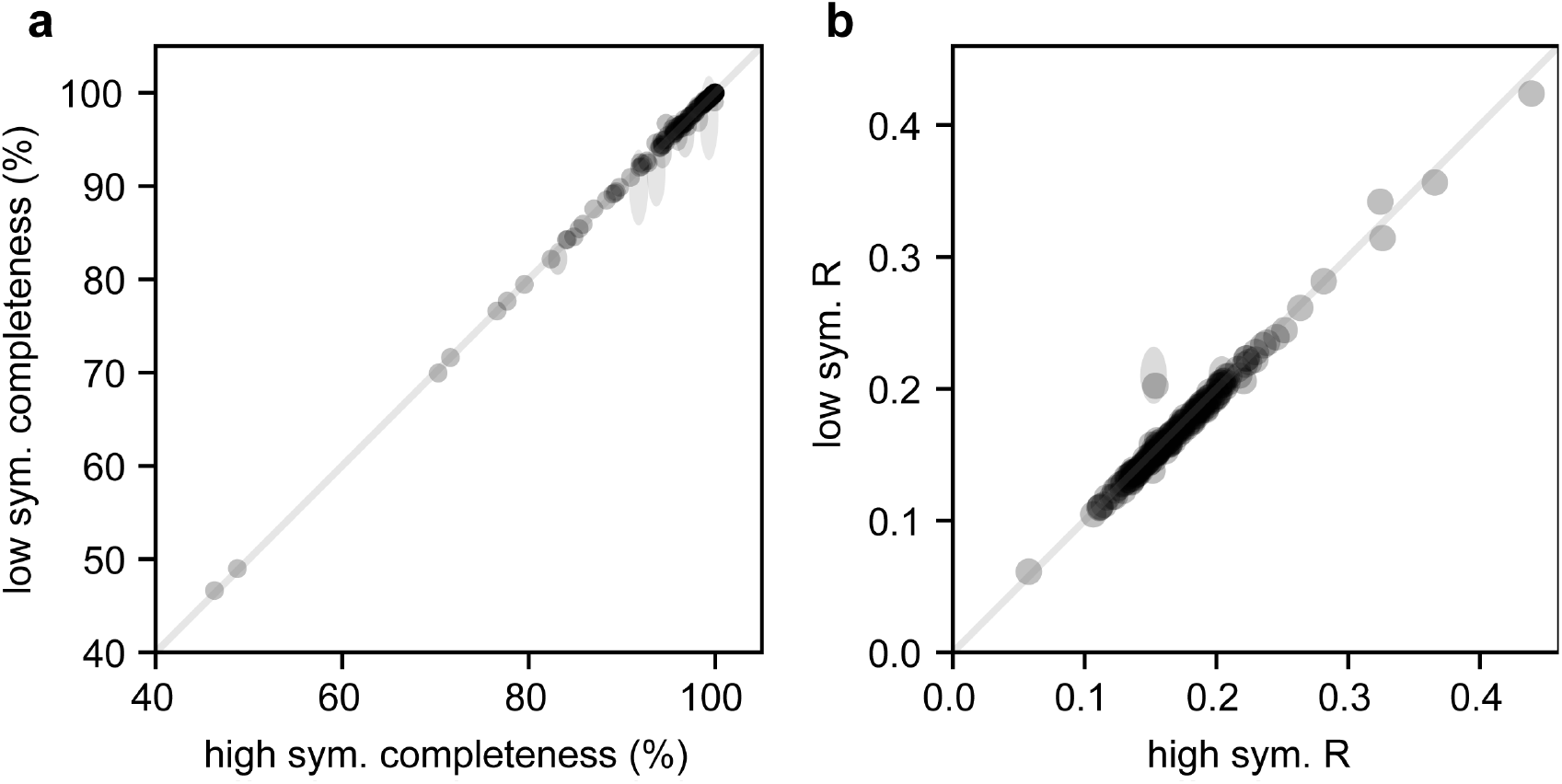
Refinement of simulated symmetry-broken data. For a given PDB entry, we plot the **a)** completeness and **b)** the R-factor of the high-symmetry data (and model), against the completeness or R-factor of the low-symmetry data (and model), respectively. R-factor is calculated after a rigid-body refinement of the high-symmetry data against the high-symmetry model, or a rigid-body refinement of the low-symmetry data against the low-symmetry model. Simulated low-symmetry data is computed by applying a simulated vectorial perturbation pointing in the specified direction and then expanding the high-symmetry data and model to the appropriate space group. Four different perturbation directions were chosen: (1,0,0), (0,1,0), (1,1,0), and (1,-1,-1). Completeness and R-factors were calculated for each low-symmetry dataset, then aggregated by mean and standard devation. For each metric, each point is centered on the mean and scaled vertically by two times the standard deviation, down to a minimum height.

## V. DISCUSSION

In this work, we formally treat symmetry breaking in crystallography from vectorial perturbations, demonstrating how to analyze both real and simulated symmetry-broken data in any Sohncke space group. We anticipate we can apply this formalism to crystallographic data in which symmetry breaking is present but not conventionally modeled. We also anticipate applying this formalism to non-Sohncke space groups for study of symmetry breaking in small molecules^44^. To our understanding, this is straightforward but not yet tested comprehensively (**Section IV D**).

Symmetry breaking is yet unmodeled in chromophore excitation experiments. Even though chromophore excitation can strongly depend on the chromophore orientation with respect to polarization direction of the incident light^45^, typical time-resolved crystallography experiments rely on light that has been depolarized using a filter. Crystal symmetry meaningfully influences the chromophore orientation, although orientation-dependent effects are typically averaged out due to processing in the high-symmetry space group. In one time-resolved crystallography experiment using linearly-polarized light stimulation, the crystal orientation was observed to affect the degree of conformational change from photochemical processes^46^. In that work, the crystal orientation varied over the serial crystallography experiment. To measure orientation-dependent effects, each image was placed into one of twelve orientation bins, then the average chromophore orientation of each bin and difference density were calculated. This analysis was possible because the average chromophore orientation in the unit cell was such that the transition dipole moment “preferred” to lie in the crystallographic *ab* plane, but this preference is not guaranteed for a general space group. Orientation anisotropy depends on both space group and the details of the studied macromolecule. By combining per-orientation binning with symmetry breaking analysis, we envision that in linearly-polarized light-stimulated crystallography, we can directly correspond the chromophore orientation to the degree of conformational change from photochemical processes. There remains one technical detail when analyzing light-stimulated experiments: the applied electric field is not a simple dipole, since a chromophore with a transition dipole moment 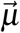 is excited the same as one with a negative 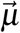. In this case, instead of preserving only symmetry operations with FSA near 1, we now preserve symmetry operations with FSA near 1 or −1 (**Section II**).

## VI. METHODS

### A. HRas purification and crystallization

For HRas dataset 1, the plasmid was generously gifted by Dr. Carla Mattos. The resulting protein was expressed and purified as described previously^50^, except that after the anion exchange column, we buffer exchanged the protein into nucleotide exchange buffer using PD Spintrap G-25 (GE) columns and used 17 units of soluble alkaline phosphatase per 10 mg of crude protein. Following nucleotide exchange, we performed size-exclusion chromatography following ref. 51. We then buffer exchanged the protein into 20 mM Tris pH 8.0, 50 mM NaCl, 5 mM MgCl_2_, 1 mM DTT, 5% glycerol, and 20 *µ*M GMPPNP and crystallized it as in ref. 52. We equilibrated sitting drops of 2 *µ*L protein and 2 *µ*L well solution against 500 uL of well solution, namely, 20-22% PEG 3350, 200 mM CaCl_2_. In some wells, crystals spontaneously grew. These crystals were used for microseeding wells without crystals.

For HRas dataset 2, the protein was generously gifted by Dr. Christine Gee, and crystallized as described previously^51^.

### B. HRas monochromatic data collection and refinement

We collected a room-temperature dataset on an HRas crystal at the National Synchrotron Light Source-II (NSLS-II) beamline 17-ID-2 at Brookhaven National Laboratory, during one beamtime allocation on February 4, 2025. We collected diffraction data at 298 K using helical data collection with a beam size at 10 x 10 *µ*m, transmission of 0.005%, exposure time of 0.05 seconds, beam energy of 12.66 keV, and over 180 images at 1° oscillation. We reduced these data using DIALS version 3.21.1 and subsequently refined a room-temperature model against these data, starting from concatenating PDB IDs 3K8Y (as conformer B) and 2RGE (as conformer A). We performed occupancy, coordinate, and anisotropic B factor refinement using `phenix.refine`^49^.

### C. Ras EF-X data collection

We collected dataset 1 on a single HRas crystal at 287 K at the BioCARS beamline 14-ID-B at Argonne National Laboratory. We collected this dataset during a beamline allocation on April 15, 2025, used the solid-state experimental apparatus^8^ and a tungsten wire with diameter 41 *µ*m for the bottom electrode. We mounted this crystal in the BioCARS environmental room at 10°C and 90+% humidity. For each *ϕ* angle, we collected a diffraction image without the electric field (“off”), then an image during a positive high-voltage pulse (“+V”), and then an image at a negative high-voltage pulse (“-V”). After the three images, we rotated the crystal around the *ϕ* axis and repeated the collection sequence. The nominal repetition rate was 1 Hz. The Laue X-ray pulses had a beam size of 50 *µ*m tall by 90 *µ*m wide, duration of 100 ps, energy of 5.5 *µ*J, and a spectrum from 1.02 - 1.11 Å and peaked at 1.04 Å, with a bandwidth of about 3%.

For the first two passes of dataset 1, we collected data at 2° *ϕ* steps at *ϕ* angles of 0 to 120° and 121 to 241°, respectively. We collected the +V and −V data 200 ns into 250 ns pulses of nominally +2 kV and −2 kV, respectively. For the third pass of dataset 1, we collected data at 1° *ϕ* steps at *ϕ* angles of 241.5-361.5°, and we collected the +V and −V data 200 ns into 250 ns pulses of nominally +3 kV and −3 kV, respectively.

We collected dataset 2 on a single HRas crystal at 277 K at the BioCARS beamline 14-ID-B at Argonne National Laboratory. We collected this dataset during a beamline allocation on August 2, 2021, used the solid-state experimental apparatus and a 41 *µ*m tungsten wire. For each *ϕ* angle, we collected a diffraction image without the electric field (“off”), then an image during a positive high-voltage pulse (“+V”). After the two images, we rotated the crystal around the *ϕ* axis and repeated the collection sequence. The Laue X-ray pulses had a beam size of 50 *µ*m tall by 90 *µ*m wide, duration of 100 ps, and a spectrum from 1.02 to 1.18 Å and peaked at 1.04 Å, with a bandwidth of about 5%.

For the first two passes of dataset 2, we collected data at 2° *ϕ* steps at *ϕ* angles of 0 to 60° and 61 to 121°, and we collected the ON data 200 ns into 250 ns pulses of nominally +3 kV, respectively. For the next three passes of dataset 2, we collected data at 1° *ϕ* steps at *ϕ* angles of 121.5 to 181.5, 181.5 to 241.75, and 252.5 to 302.25°, and we collected the ON data 200 ns into 250 ns pulses of nominally +3 kV, respectively.

### D. HRas EF-X data processing

We processed HRas datasets 1 and 2 using the workflow in Figure 2. We indexed and integrated the data in Precognition (Renz Research, Inc.) in spacegroup R32:H, then used regroup to compute the field-symmetry alignment for each symmetry operation in each dataset, the reduced-symmetry space group, and the P1 unmerged reflections. Then, we scaled and merged the data using Careless 0.5.3 with 3 half-dataset repeats for cross-validation. We performed ablations by using default **h**_**obs**_ as metadata instead of the high-symmetry **h**_**obs**_, or by avoiding reindexing low-symmetry MTZ files prior to scaling, or by removing the `--double_wilson_reindexing_ops` flag, or combinations of these. All ablations correspond to details discussed in III E. We used `reciprocalspaceship` to compute CC_sym_s on these half-dataset repeats. Starting from a room-temperature model of HRas, we constructed a model of the P1 asymmetric unit by first making a template model through molecular replacement of 6 copies of the high-symmetry ASU, then generating another model built by symmetry-expanding the high-symmetry ASU using the PyMOL psico module (available at https://github.com/speleo3/PyMOL-psico). The latter model was then aligned to the template model to yield an initial low-symmetry model.

### E. PDZ3 EF-X data processing

PDZ3 EF-X data included 264 images at both off and 200ns timepoints with 4 kV of applied field. We processed these data using the workflow in Figure 2. We indexed and integrated the data in Precognition in spacegroup P4_1_32, then computed the field-symmetry alignment of each dataset, the reduced-symmetry space group, and the R3:H unmerged reflections using regroup. Then, we scaled and merged the data using Careless 0.5.3. We used the double-Wilson prior to impose the expected correlation between the OFF P4_1_32 data and the OFF R3:H data, as well as a correlation between the ON P4_1_32 data and the OFF R3:H data. Thus, the OFF data were double-counted. In previous work, we found that this does not cause issues^40^. Careless produced three merged datasets: an OFF, P4_1_32 dataset, an OFF, R3:H dataset, and an ON, R3:H dataset. Careless also provided 3 half-dataset repeats for cross-validation. Such half-datasets were merged independently. We used `reciprocalspaceship` to compute CC_sym_s on these half-dataset repeats, and then make internal difference maps. One internal difference map was computed per broken symmetry operation, where structure factor amplitude differences are computed as

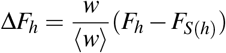

with a symmetry operation *S* transforming the Miller indices *h* and the weights *w* computed as

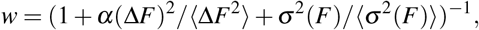

with ⟨⟩ denoting the mean of *F*, the structure factor amplitude, or *σ F*, the error on the structure factor amplitude, and *α* is a weight term chosen to be 0.05.

The internal difference maps were phased with an initial low-symmetry model that had been rigid-body refined against the OFF structure factor amplitudes.

We also used `reciprocalspaceship` to compute weighted extrapolated structure factor amplitudes between the low-symmetry off data and on data following the formula: 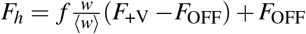, where *w* is defined as before, and *f* is the extrapolation factor. To control for refinement artifacts, we also computed extrapolated structure factor amplitudes between the R3:H OFF data and R3-expanded P4_1_32 OFF data.

Starting from a room-temperature model of PDZ3, we constructed a model of the R3:H asymmetric unit empirically by molecular replacement of 8 copies of the high-symmetry asymmetric unit following the procedure described in **Section VI D**. Then, we performed automated refinement of the coordinates and occupancies using phenix.refine. We then extracted coordinates of the refined model and performed singular value decomposition (SVD) on the coordinate displacements between the ON and OFF data, across all 8 molecules in the asymmetric unit, following ref. 16. Also as in ref. 16, we chose the extrapolation factor in **Figure 4f** by determining a plateau in a coordinate-based metric. Specifically, we noted a quantity that varied near-monotonically as a function of the extrapolation factor: the correlation coefficient of the singular vector loadings between ASU molecules that were oppositely oriented to the electric field (**Figure 4d**, **S5**). We found a plateau at extrapolation factor 12 with a correlation coefficient near −1 for both SV1 and SV2 loadings. We did not find the same trend when we refined models against the control extrapolation data. In fact, we found that the correlation coefficient converged to 1. This likely indicates that the OFF-OFF differences contained noise that reduced the strength of X-ray data restraints during refinement relative to the strength of geometric restraints. On the other hand, the OFF-ON differences were most likely due to the electric field and opposite in chains oppositely oriented to the electric field.

### F. Automated symmetry reduction

To test our symmetry-breaking method, we wrote an automated routine for low-symmetry model generation, and tested it on simulated symmetry breaking on data from the Protein Data Bank. Our test set was a random set of 200 protein, DNA, and peptide entries in the PDB across 77 spacegroups (including all chiral space groups and the centrosymmetric space groups P12_1_/c1, P12_1_/n1, 12_1_/n1, C12/n1, P-3, H-3, and P-1). We simulated applying a vectorial perturbation normal to the (1,0,0), (0,1,0), (1,1,0), and (1,-1,-1) facets by using `regroup` to generate low-symmetry space groups, leading to a total of 800 low-symmetry maps and models. We then used `regroup.low_sym` to expand the high-symmetry MTZ files to low symmetry. Rather than add noise, we simply generated reflections with duplicate structure factor amplitudes. Then, we generated low-symmetry models by first enumerating the broken symmetry operations of the high-symmetry spacegroup, and then iterating over equivalent broken symmetry operations and centering operations to find the most compact arrangement of the symmetry mates of the high-symmetry ASU. This script is implemented as `regroup.change_basis_pdb`. We performed an affine transformation of the fractional coordinates of these atomic coordinates using the `regroup` change-of-basis operation in the setting of the low-symmetry space group. Finally, we stripped waters and ions on special positions and performed rigid-body refinement without R_free_ flags. We filtered each PHENIX run for PHENIX failure modes, e.g., if there were multiple models in one model file, atoms in rigid groups on special positions, or unknown atom types. Seven such runs failed. In two other refinement failures, PHENIX chose an improper origin for the high-symmetry model and so the high-symmetry R-factors were very large (>0.6). In one of these cases, the data were also less than 10% complete according to PHENIX, even though in reality, the data were 93% complete^53^. This indicated that out-of-the-box, PHENIX could not handle data of this particular spacegroup (Pbca). The one remaining refinement failure involved a virus model that timed out after four hours. In still other cases, to get PHENIX to run, we ignored clashes and/or removed ligands. This resulted in 190 remaining datasets for testing.

## ACKNOWLEDGMENTS

We thank Drs. Christine Gee and John Kuriyan for the HRas protein, Dr. Carla Mattos for the HRas plasmid, and Yujiao Wu, Michael Socolich, and Dr. Rama Ranganathan for the PDZ3 EF-X dataset. We thank Jeffrey Chang for feedback on the manuscript. We thank members of the Hekstra lab, particularly Dr. Dennis Brookner, for collecting the HRas dataset 2 and for discussion. We thank Dr. Shao-Liang Zheng for support with the room-temperature rotating-anode crystallography that was preliminary to EF-X. This research used resources 17-ID-2 of the National Synchrotron Light Source II (NSLS-II), a U.S. Department of Energy (DOE) Office of Science User Facility operated for the DOE Office of Science by Brookhaven National Laboratory under Contract No. DE-SC0012704. We thank the staff of the Center for BioMolecular Structure (CBMS) at the NSLS-II, especially Dr. Wuxian Shi, Dr. Alexei Soares, and Dr. Martin Fuchs, for supporting our room-temperature crystallography experiments. The CBMS is primarily supported by the National Institutes of Health (NIH), National Institute of General Medical Sciences (NIGMS) through a Center Core P30 Grant (P30GM133893), and by the DOE Office of Biological and Environmental Research (KP1605010). Use of the 14-ID-B beamline was also supported by the NIGMS of the NIH under grant number P41 GM118217. The time-resolved instrumentation at the 14-ID-B beamline was funded in part through a collaboration with Philip Anfinrud (NIH/National Institute of Diabetes and Digestive and Kidney Diseases). This work was also supported by a NIH grant DP2-GM141000 (to D.R.H.), National Science Foundation Graduate Research Fellowship grant DGE2140743 (to H.K.W.), Burroughs Wellcome Fund Career Award at the Scientific Interface (to K.M.D.), and a National Science Foundation Graduate Research Fellowship grant DGE1745303 (to J.B.G.).

## DATA AVAILABILITY STATEMENT

The data that support the findings of this study are openly available in a Zenodo deposition at https://zenodo.org/records/20818958. Regroup is available at https://github.com/Hekstra-Lab/regroup. Refined models have been deposited in the Protein Data Bank with IDs 36OR, 36OU, XXXX, and XXXX.

## VII. SUPPLEMENTARY INFORMATION

### A. Closed form for the FSA

We stated in section II B that if we rotate a field’s unit vector 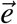 by *ϕ* degrees, and the unit vector is *θ* degrees away from the rotation axis, then the FSA is cos^2^ *θ* + sin^2^ *θ* cos *ϕ*. To see why this is true, let’s work in Euclidean space and break the unit vector 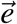 into the components parallel 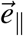 and perpendicular 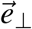 to the symmetry axis, so that 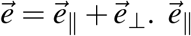 has length cos *θ* and 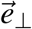 has length sin *θ*.

Now call the rotated vector 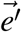.

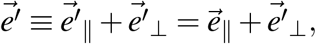

as 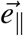 is parallel to the rotation axis and unchanged by rotation. Now note 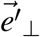 has been rotated *ϕ* away from 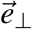. So

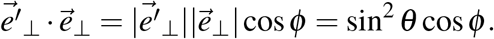

Finally, the FSA (in Euclidean space) is

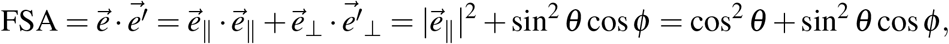

as desired.

**TABLE S1:**
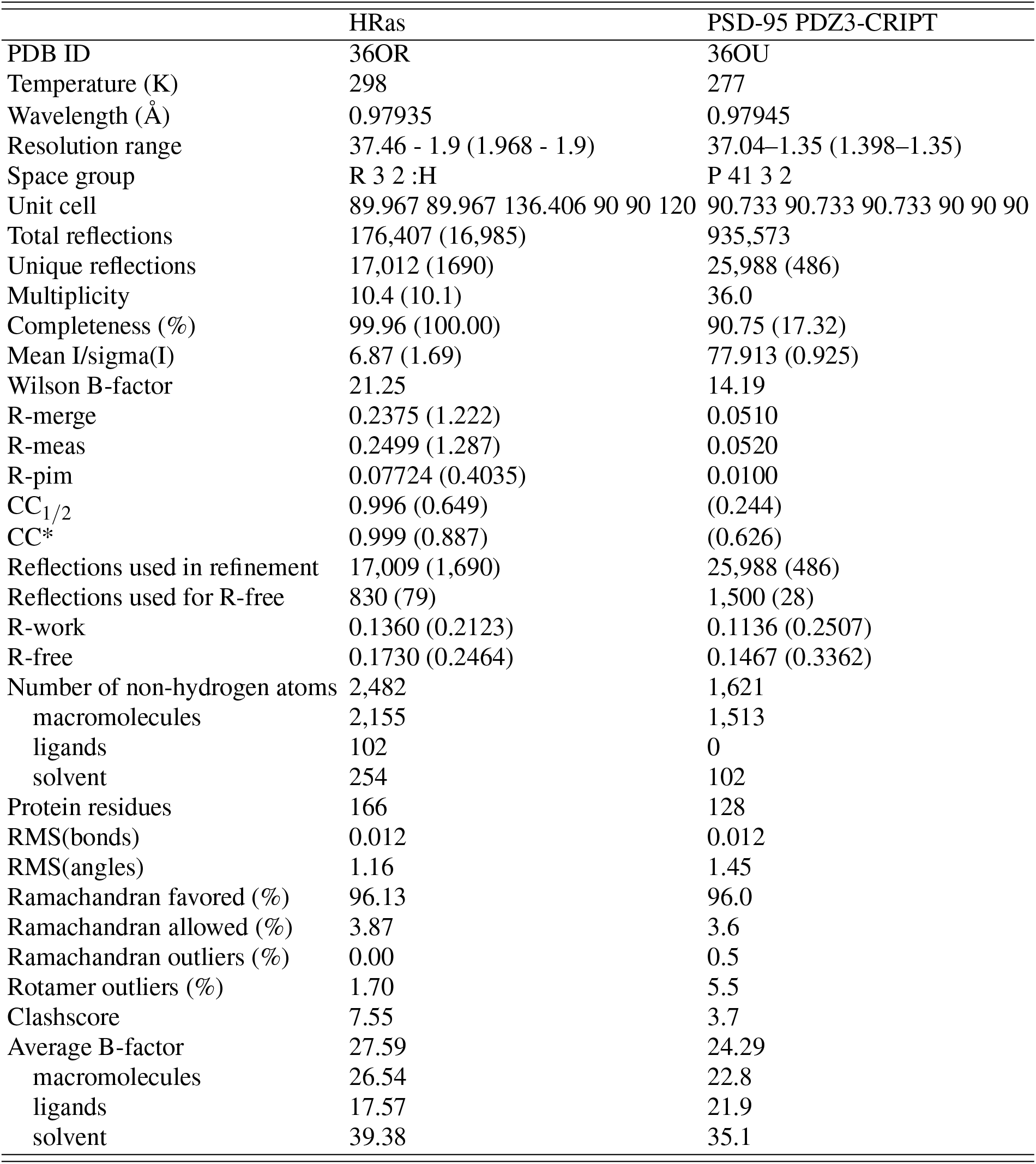
Summary statistics for room-temperature monochromatic datasets in this manuscript.

**TABLE S2:** Data reduction statistics for PDZ3 EF-X data. Statistics were computed based on output from Careless.

| Dataset | OFF | OFF (low sym.) | ON |
| --- | --- | --- | --- |
| No. of images | 264 | 264 | 264 |
| Space group | P4<sub>1</sub>32 | R3:H | R3:H |
| Cell dim. |  |  |  |
| a (Å) | 90.5 | 127.986 | 127.986 |
| b (Å) | 90.5 | 127.986 | 127.986 |
| c (Å) | 90.5 | 156.751 | 156.751 |
| $\alpha$ (°) | 90 | 90 | 90 |
| $\beta$ (°) | 90 | 90 | 90 |
| $\gamma$ (°) | 90 | 120 | 120 |
| Total obs. | 1,199,649 (91,316) | 1,199,649 (91316) | 1,193,904 (90384) |
| Unique obs. | 10,506 (1,043) | 75,417 (7,729) | 75,412 (7,724) |
| Resolution | 63.99 - 1.90 (1.97 - 1.90) | 63.99 - 1.90 (1.97 - 1.90) | 63.99 - 1.90 (1.97 - 1.90) |
| Multiplicity | 114.2 (87.6) | 15.91 (11.8) | 15.8 (11.7) |
| Completeness | 1.0 (1.0) | 0.999 (1.0) | 0.999 (0.998) |
| Mean I/$\sigma$ I | 773 (87) | 275 (26) | 272 (26) |
| CC<sub>1/2</sub> | 0.998 (0.994) | 0.999 (0.992) | 0.999 (0.991) |

**TABLE S3:**
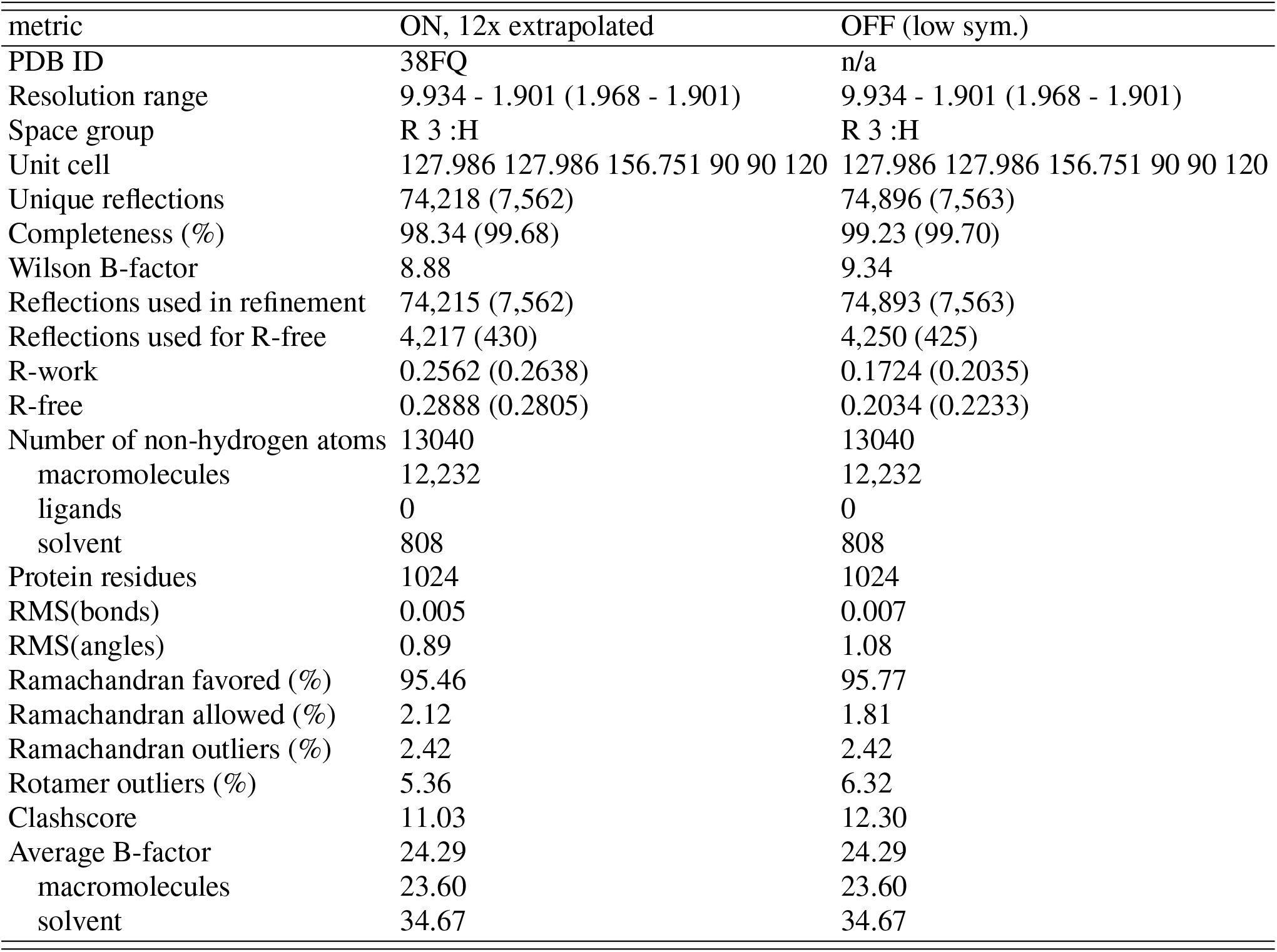
Refinement statistics for PDZ3 EF-X data.

| metric | ON, 12x extrapolated | OFF (low sym.) |
| --- | --- | --- |
| PDB ID | 38FQ | n/a |
| Resolution range | 9.934 - 1.901 (1.968 - 1.901) | 9.934 - 1.901 (1.968 - 1.901) |
| Space group | R 3 :H | R 3 :H |
| Unit cell | 127.986 127.986 156.751 90 90 120 | 127.986 127.986 156.751 90 90 120 |
| Unique reflections | 74,218 (7,562) | 74,896 (7,563) |
| Completeness (%) | 98.34 (99.68) | 99.23 (99.70) |
| Wilson B-factor | 8.88 | 9.34 |
| Reflections used in refinement | 74,215 (7,562) | 74,893 (7,563) |
| Reflections used for R-free | 4,217 (430) | 4,250 (425) |
| R-work | 0.2562 (0.2638) | 0.1724 (0.2035) |
| R-free | 0.2888 (0.2805) | 0.2034 (0.2233) |
| Number of non-hydrogen atoms | 13040 | 13040 |
| macromolecules | 12,232 | 12,232 |
| ligands | 0 | 0 |
| solvent | 808 | 808 |
| Protein residues | 1024 | 1024 |
| RMS(bonds) | 0.005 | 0.007 |
| RMS(angles) | 0.89 | 1.08 |
| Ramachandran favored (%) | 95.46 | 95.77 |
| Ramachandran allowed (%) | 2.12 | 1.81 |
| Ramachandran outliers (%) | 2.42 | 2.42 |
| Rotamer outliers (%) | 5.36 | 6.32 |
| Clashscore | 11.03 | 12.30 |
| Average B-factor | 24.29 | 24.29 |
| macromolecules | 23.60 | 23.60 |
| solvent | 34.67 | 34.67 |

**TABLE S4:** Data reduction statistics for Ras EF-X dataset 1. Statistics were computed based on output from Careless.

| Dataset | OFF | OFF (low sym.) | +V | -V |
| --- | --- | --- | --- | --- |
| No. of images | 243 | 243 | 243 | 243 |
| Space group | R32:H | P1 | P1 | P1 |
| Cell dim. (Å) |  |  |  |  |
| a (Å) | 89.959 | 68.878 | 68.878 | 68.878 |
| b (Å) | 89.959 | 68.878 | 68.878 | 68.878 |
| c (Å) | 135.718 | 68.878 | 68.878 | 68.878 |
| $\alpha$ (°) | 90 | 81.542 | 81.542 | 81.542 |
| $\beta$ (°) | 90 | 81.542 | 81.542 | 81.542 |
| $\gamma$ (°) | 120 | 81.542 | 81.542 | 81.542 |
| Total obs. | 282,648 (21,511) | 282,648 (21,511) | 261,630 (18,727) | 261,473 (18,703) |
| Unique obs. | 16,917 (1,723) | 89018 (8,522) | 87,493 (8,120) | 87,493 (8,120) |
| Resolution | 67.57 - 1.90 (1.97 - 1.90) | 67.57 - 1.90 (1.97 - 1.90) | 67.57 - 1.90 (1.97 - 1.90) | 67.57 - 1.90 (1.97 - 1.90) |
| Multiplicity | 16.7 (12.5) | 3.17 (2.52) | 2.99 (2.31) | 2.99 (2.31) |
| Completeness | 0.999 (1.0) | 0.919 (0.852) | 0.903 (0.814) | 0.904 (0.818) |
| Mean I/$\sigma$ I | 124.8 (55.4) | 43.0 (14.7) | 41.1 (14.6) | 41.1 (14.6) |
| CC<sub>1/2</sub> | 0.991 (0.965) | 0.988 (0.971) | 0.987 (0.973) | 0.987 (0.973) |

**TABLE S5:** Data reduction statistics for Ras EF-X dataset 2. Statistics were computed based on output from Careless.

| Dataset | OFF | OFF (low sym.) | ON |
| --- | --- | --- | --- |
| No. of images | 184 | 184 | 184 |
| Space group | R32:H | P1 | P1 |
| Cell dim. (Å) |  |  |  |
| a (Å) | 89.959 | 69.108 | 69.108 |
| b (Å) | 89.959 | 69.108 | 69.108 |
| c (Å) | 136.766 | 69.108 | 69.108 |
| $\alpha$ (°) | 90 | 81.213 | 81.213 |
| $\beta$ (°) | 90 | 81.213 | 81.213 |
| $\gamma$ (°) | 120 | 81.213 | 81.213 |
| Total obs. | 214,646 (10,934) | 214,646 (10,934) | 207,091 (10,000) |
| Unique obs. | 14,324 (2,008) | 68,048 (6,265) | 66,794 (7,724) |
| Resolution | 37.46 - 1.99 (2.12 - 1.99) | 37.46 - 1.99 (2.12 - 1.99) | 37.46 - 1.99 (2.12 - 1.99) |
| Multiplicity | 15.0 (5.45) | 3.15 (1.75) | 3.10 (1.70) |
| Completeness | 1.0 (1.0) | 0.999 (1.0) | 0.999 (0.998) |
| Mean I/σI | 158.3 (49.8) | 61.9 (17.0) | 59.2 (16.8) |
| CC<sub>(1/2)</sub> | 0.989 (0.969) | 0.994 (0.978) | 0.994 (0.979) |

**FIG. S1:**
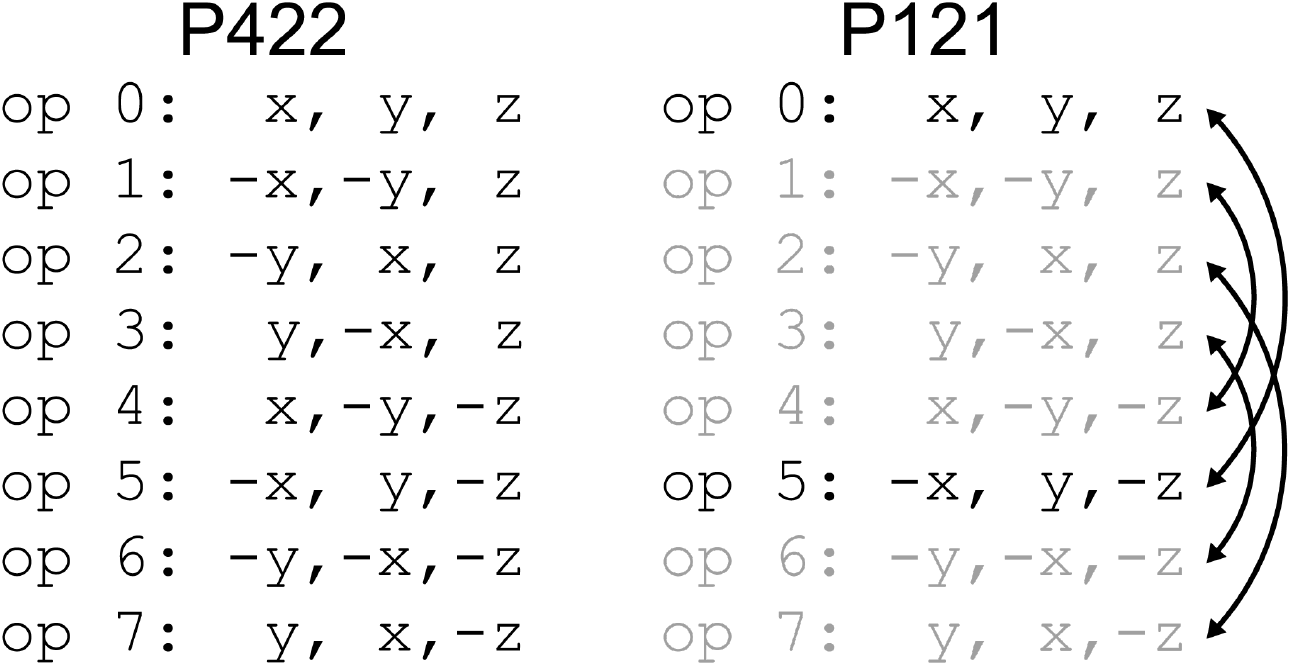
Lists of symmetry operations of P422 and its subgroup, P121. Broken rotation symmetries of P422 are greyed out in the list for P121. Symmetry-equivalent rotations in P121 are indicated by arrows. Note that rotations are equivalent if they are identical up to the preserved symmetry in P121.

**FIG. S2:**
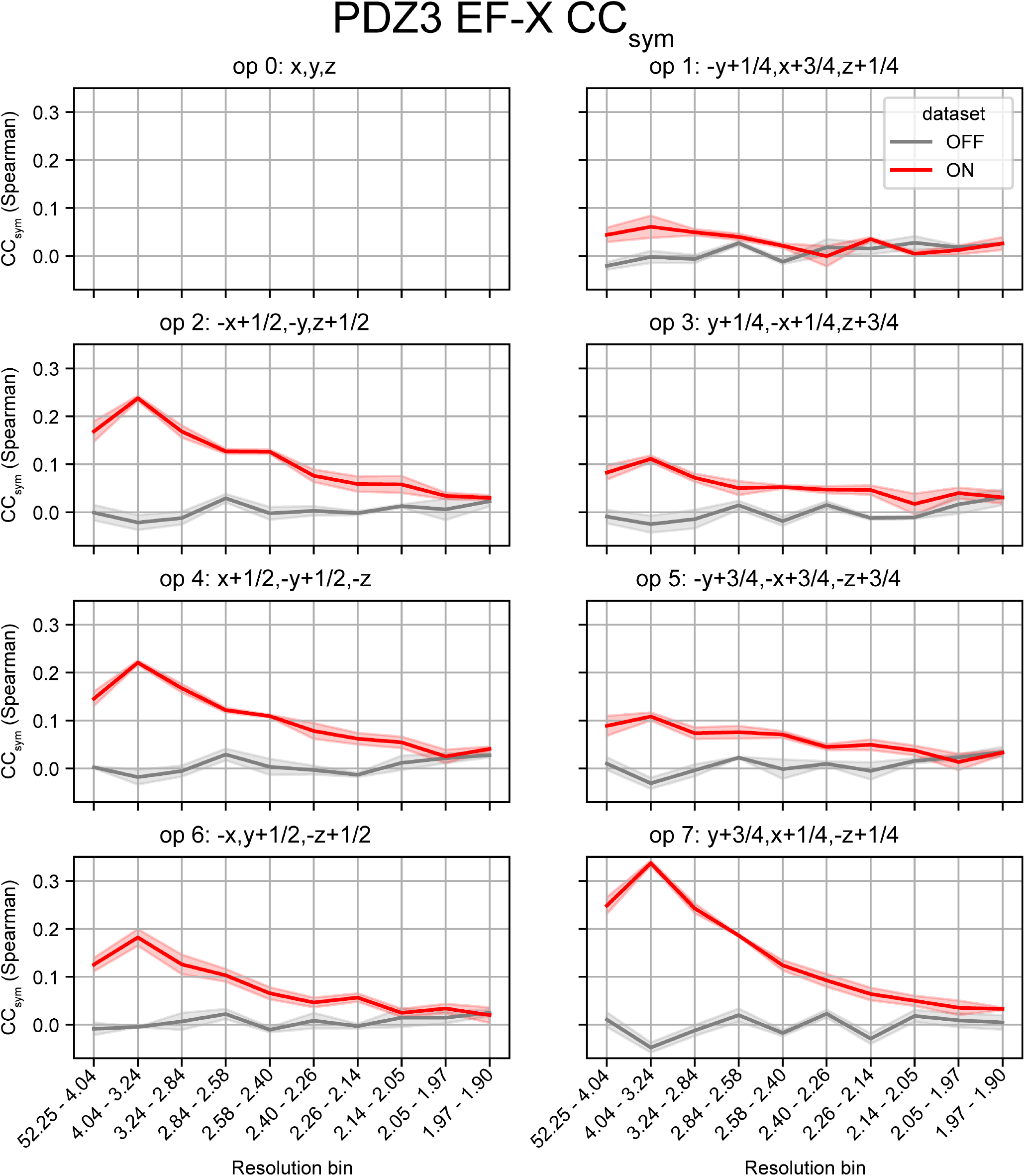
CC_sym_s of PDZ3 EF-X data. Shown are all CC_sym_s for each coset over 10 resolution bins. Red trace indicates the CC_sym_ of the PDZ3 EF-X ON dataset in R3:H, and gray trace indicates that of the PDZ3 EF-X off dataset in R3:H. The gray and red shaded regions are the standard deviation of the CC_sym_ at each resolution bin across half-dataset splits. The CC_sym_ is 0 for x,y,z by definition and is not shown.

**FIG. S3:**
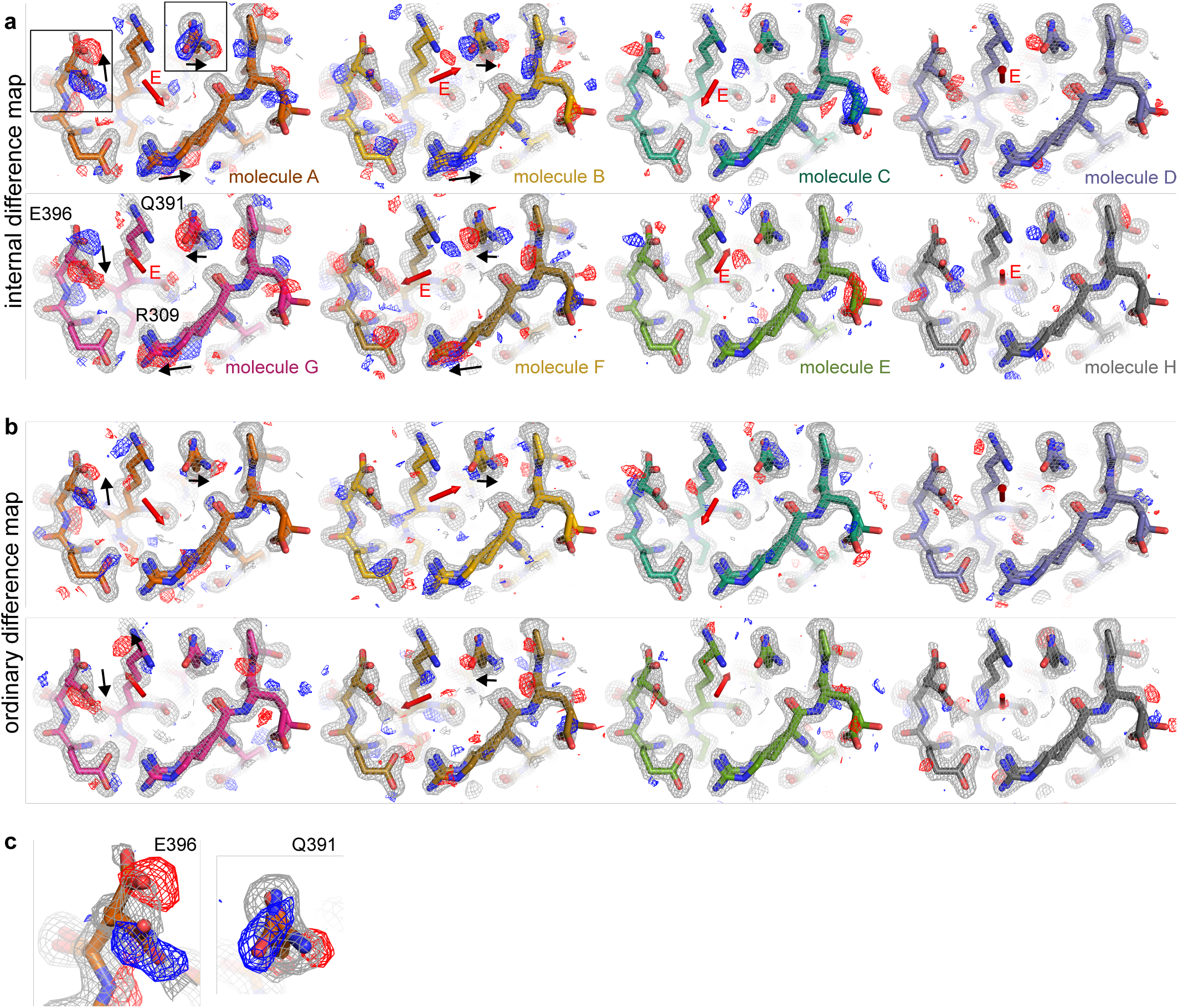
PDZ3 EF-X difference maps. **a)** internal difference maps and **b)** ordinary difference maps from the PDZ3 EF-X experiment. Insets in molecule A’s internal difference map are shown in **c)**. In all images, the OFF model is shown as sticks colored as in **Figure 4**. The F_*o*_ map is shown as a gray mesh, contoured at 1.2*σ* and carved to 1.7 Å, and the internal difference map is shown as a red and blue mesh, carved at 1.7 Å and contoured to 2.5*σ* and −2.5*σ*, respectively. Black arrows denote occupancy changes between alternate conformations that are evident in the difference maps.

**FIG. S4:**
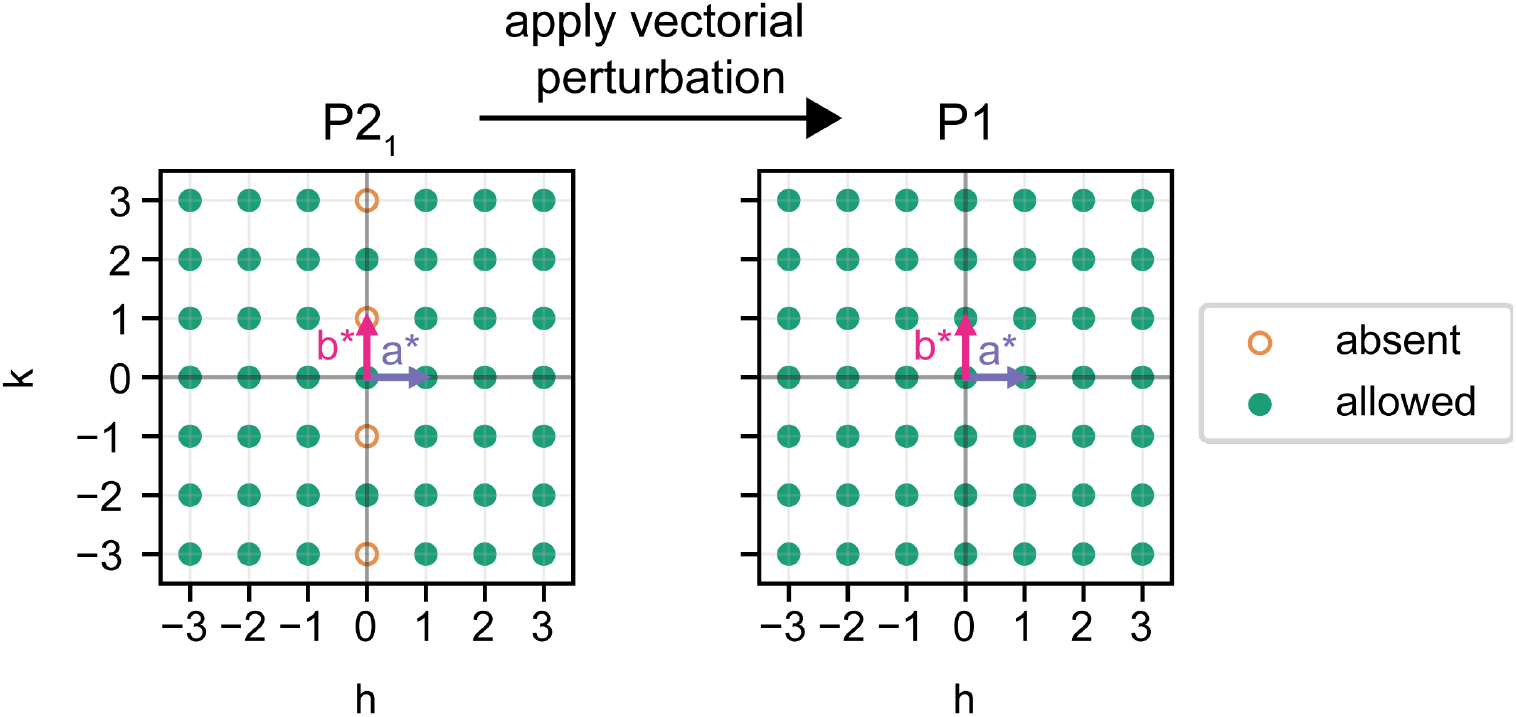
Reappearance of systematic absences from symmetry breaking in P2_1_. Plot of the reciprocal grid along *h* and *k* in P2_1_ and in P1, showing reappearance of P2_1_ systematic absences upon symmetry breaking. Absent reflections are shown as circles with orange dashed borders. Allowed reflections are shown as green disks.

**FIG. S5:**
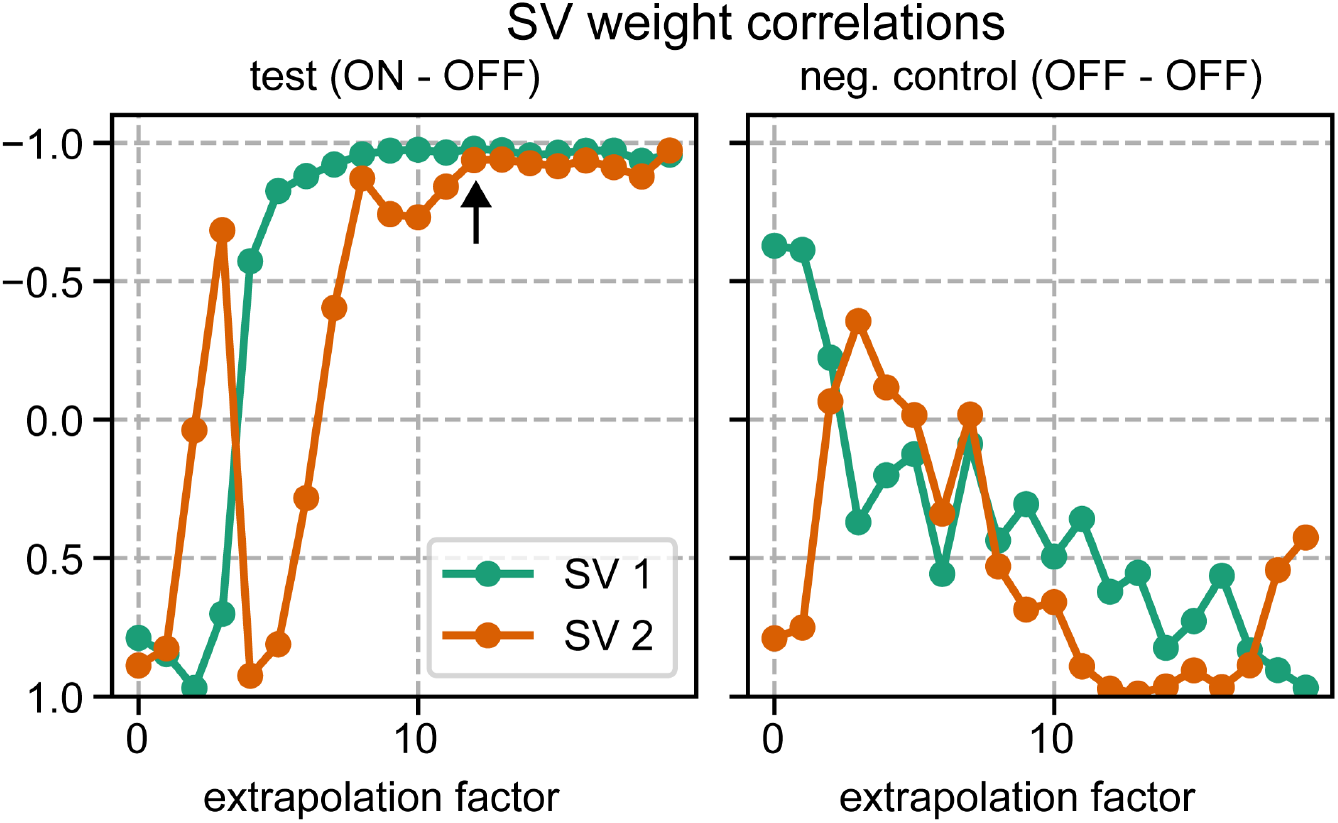
The correlation between the ABCD and GFEH molecule SVD loadings, and the dependence of this correlation on extrapolation factor. Plot of the correlation between the PDZ3 ABCD and GFEH molecule SVD loadings, for the first two singular vectors (SVs), as in **Figure 4f**. Note that the *y* axis has been inverted. When the true differences between ON and OFF data are extrapolated, this correlation becomes more negative with increasing extrapolation factor. When the spurious differences between OFF,R3:H and OFF,P4132 data are extrapolated, this correlation becomes more positive with increasing extrapolation factor. This indicates that extrapolation amplifies oppositely-oriented symmetry-breaking signal in the ON data. Black arrow indicates the chosen extrapolation factor, where both the SV1 and SV2 correlations have plateaued.

**FIG. S6:**
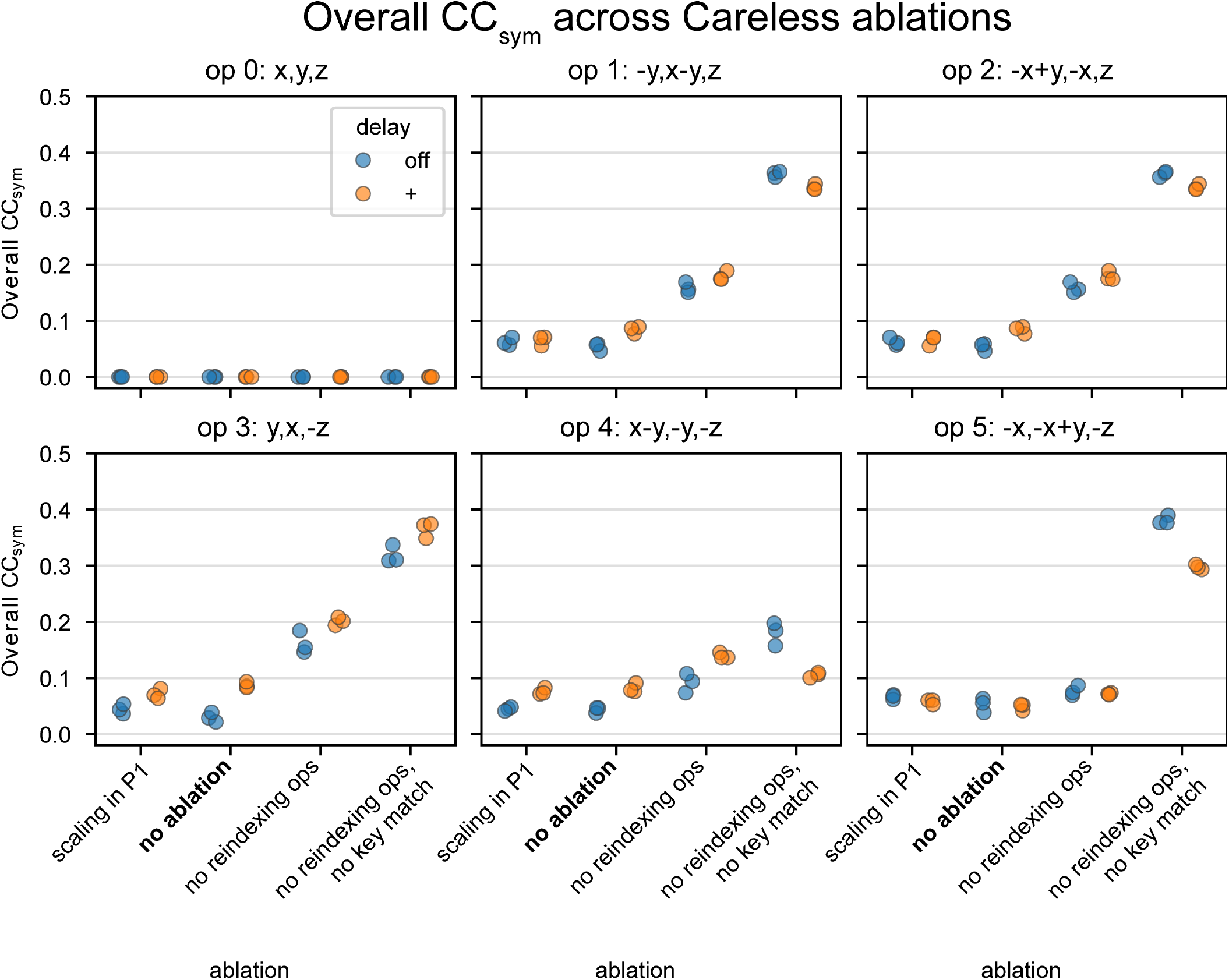
CC_sym_ of Ras EF-X dataset 2 for various Careless metadata settings. Shown are all CC_sym_ for each coset over all reflections. The blue circles represent the overall CC_sym_ for the OFF dataset and the orange circles represent the overall CC_sym_ for the ON (+) dataset. For a given coset, the blue and orange circles are repeated four times, one for each Careless metadata setting as in **Figure 5f**. This plot indicates that essential to good scaling is handling basis change and observed Miller indices correctly. Poor scaling can lead to residual systematic errors and inflated CC_sym_ even in the OFF data, as discussed in **Section**.

## References

1 A. M. Wolff, E. Nango, I. D. Young, A. S. Brewster, M. Kubo, T. Nomura, M. Sugahara, S. Owada, B. A. Barad, K. Ito, et al., Nature chemistry 15, 1549 (2023).

2 P. Mehrabi, E. C. Schulz, R. Dsouza, H. M. Müller-Werkmeister, F. Tellkamp, R. D. Miller, and E. F. Pai, Science 365, 1167 (2019).

3 T. Gruhl, T. Weinert, M. J. Rodrigues, C. J. Milne, G. Ortolani, K. Nass, E. Nango, S. Sen, P. J. Johnson, C. Cirelli, et al., Nature 615, 939 (2023).

4 T. R. Barends, L. Foucar, A. Ardevol, K. Nass, A. Aquila, S. Botha, R. B. Doak, K. Falahati, E. Hartmann, M. Hilpert, et al., Science 350, 445 (2015).

5 T. R. Barends, A. Gorel, S. Bhattacharyya, G. Schirò, C. Bacellar, C. Cirelli, J.-P. Colletier, L. Foucar, M. L. Grünbein, E. Hartmann, et al., Nature 626, 905 (2024).

6 V. Šrajer, Z. Ren, T.-Y. Teng, M. Schmidt, T. Ursby, D. Bourgeois, C. Pradervand, W. Schildkamp, M. Wulff, and K. Moffat, Biochemistry 40, 13802 (2001).

7 S. Mous, G. Gotthard, D. Ehrenberg, S. Sen, T. Weinert, P. J. Johnson, D. James, K. Nass, A. Furrer, D. Kekilli, et al., Science 375, 845 (2022).

8 B. Lee, K. I. White, M. Socolich, M. A. Klureza, R. Henning, V. Srajer, R. Ranganathan, and D. R. Hekstra, Cell 188, 77 (2025).

9 J.-H. Yun, X. Li, J. Yue, J.-H. Park, Z. Jin, C. Li, H. Hu, Y. Shi, S. Pandey, S. Carbajo, et al., Proceedings of the National Academy of Sciences 118, e2020486118 (2021).

10 N.-E. Christou, V. Apostolopoulou, D. V. Melo, M. Ruppert, A. Fadini, A. Henkel, J. Sprenger, D. Oberthuer, S. Günther, A. Pateras, et al., Science 382, 1015 (2023).

11 D. E. Brookner, Evolutionary Motif Swapping of Human Dihydrofolate Reductase Rewires the Enzymatic Cycle, Ph.D. thesis, Harvard University (2025).

12 H. Yamada, T. Nagae, and N. Watanabe, Biological Crystallography 71, 742 (2015).

13 A. Svanidze, H. Huth, S. Lushnikov, S. Kojima, and C. Schick, Applied Physics Letters 95 (2009).

14 S. Klingl, M. Scherer, T. Stamminger, and Y. A. Muller, Biological Crystallography 71, 1493 (2015).

15 D. R. Hekstra, K. I. White, M. A. Socolich, R. W. Henning, V. Šrajer, and R. Ranganathan, Nature 540, 400 (2016).

16 J. B. Greisman, K. M. Dalton, D. E. Brookner, M. A. Klureza, C. J. Sheehan, I.-S. Kim, R. W. Henning, S. Russi, and D. R. Hekstra, Proceedings of the National Academy of Sciences 121, e2313192121 (2024).

17 M. L. Grünbein, A. Gorel, L. Foucar, S. Carbajo, W. Colocho, S. Gilevich, E. Hartmann, M. Hilpert, M. Hunter, M. Kloos, et al., Nature communications 12, 1672 (2021).

18 I. V. Lundholm, H. Rodilla, W. Y. Wahlgren, A. Duelli, G. Bourenkov, J. Vukusic, R. Friedman, J. Stake, T. Schneider, and G. Katona, Structural Dynamics 2 (2015).

19 D. R. Hekstra, Annual review of biophysics 52, 255 (2023).

20 Y. A. Izyumov and V. N. Syromyatnikov, Phase transitions and crystal symmetry, Vol. 38 (Springer Science & Business Media, 2012).

21 G. de la Flor, M. I. Aroyo, I. Gimondi, S. C. Ward, K. Momma, R. M. Hanson, and L. Suescun, Applied Crystallography 57 (2024).

22 H. T. Stokes, B. J. Campbell, and D. M. Hatch, Foundations of Crystallography 63, 365 (2007).

23 U. Müller, Symmetry relationships between crystal structures: applications of crystallographic group theory in crystal chemistry, Vol. 18 (OUP Oxford, 2013).

24 P. H. Zwart, R. W. Grosse-Kunstleve, A. A. Lebedev, G. N. Murshudov, and P. D. Adams, Biological Crystallography 64, (2008).

25 B. K. Poon, R. W. Grosse-Kunstleve, P. H. Zwart, and N. K. Sauter, Biological Crystallography 66, 503 (2010).

26 A. A. Lebedev and M. N. Isupov, Biological Crystallography 70, 2430 (2014).

27 B. Rupp, Biomolecular crystallography: principles, practice, and application to structural biology (Garland Science, 2009).

28 M. Artin, Algebra, 2nd ed. (Pearson Prentice Hall, 2011).

29 T. Hahn, ed., International Tables for Crystallography, Volume A: Space-Group Symmetry, 5th ed. (Springer, Dordrecht, 2005).

30 H. T. Stokes, S. v. Orden, and B. J. Campbell, Applied Crystallography 49, 1849 (2016).

31 M. Wojdyr, Journal of Open Source Software 7, 4200 (2022).

32 R. W. Grosse-Kunstleve, N. K. Sauter, N. W. Moriarty, and P. D. Adams, Applied Crystallography 35, 126 (2002).

33 R. Grosse-Kunstleve, Foundations of Crystallography 55, 383 (1999).

34 J. B. Greisman, K. M. Dalton, and D. R. Hekstra, Applied Crystallography 54, 1521 (2021).

35 G. Winter, D. G. Waterman, J. M. Parkhurst, A. S. Brewster, R. J. Gildea, M. Gerstel, L. Fuentes-Montero, M. Vollmar, T. Michels-Clark, I. D. Young, et al., Biological Crystallography 74, 85 (2018).

36 J. Helliwell, J. Habash, D. Cruickshank, M. Harding, T. Greenhough, J. Campbell, I. Clifton, M. Elder, P. Machin, M. Papiz, et al., Applied Crystallography 22, 483 (1989).

37 K. M. Dalton, J. B. Greisman, and D. R. Hekstra, Nature Communications 13, 7764 (2022).

38 L. A. Aldama, K. M. Dalton, and D. R. Hekstra, Biological Crystallography 79, 796 (2023).

39 K. A. Zielinski, C. Dolamore, H. K. Wang, R. W. Henning, M. A. Wilson, L. Pollack, V. Srajer, D. R. Hekstra, and K. M. Dalton, Structural Dynamics 11 (2024).

40 D. R. Hekstra, H. K. Wang, M. A. Klureza, J. B. Greisman, and K. M. Dalton, Science Advances 11, eadj2921 (2025).

41 P. Evans, Biological crystallography 62, 72 (2006).

42 B. A. Katz, Journal of molecular biology 274, 776 (1997).

43 B. A. Katz, R. Mackman, C. Luong, K. Radika, A. Martelli, P. A. Sprengeler, J. Wang, H. Chan, and L. Wong, Chemistry & biology 7, 299 (2000).

44 L. K. Saunders, H.-M. Yeung, M. R. Warren, P. Smith, S. Gurney, S. F. Dodsworth, I. J. Vitorica-Yrezabal, A. Wilcox, P. V. Hathaway, G. Preece, et al., Applied Crystallography 54, 1349 (2021).

45 K. Ng, E. D. Getzoff, and K. Moffat, Biochemistry 34, 879 (1995).

46 S. Perrett, A. Fadini, C. D. Hutchison, H. Rycroft, S. Bhattacharya, D. Morozov, S. Muniyappan, T. N. Malla, D. Khakhulin, O. Turkot, et al., ChemRxiv (2025).

47 D. A. Doyle, A. Lee, J. Lewis, E. Kim, M. Sheng, and R. MacKinnon, Cell 85, 1067 (1996).

48 R. A. Hewitt, K. M. Dalton, D. A. Mendez, H. K. Wang, M. A. Klureza, D. E. Brookner, J. B. Greisman, D. McDonagh, V. Šrajer, N. K. Sauter, et al., Structural Dynamics 11 (2024).

49 P. D. Adams, P. V. Afonine, G. Bunkóczi, V. B. Chen, I. W. Davis, N. Echols, J. J. Headd, L.-W. Hung, G. J. Kapral, R. W. Grosse-Kunstleve, et al., Biological crystallography 66, 213 (2010).

50 C. W. Johnson, G. Buhrman, P. Y. Ting, J. Colicelli, and C. Mattos, Data in brief 6, 423 (2016).

51 P. Bandaru, N. H. Shah, M. Bhattacharyya, J. P. Barton, Y. Kondo, J. C. Cofsky, C. L. Gee, A. K. Chakraborty, T. Kortemme, R. Ranganathan, and J. Kuriyan, eLife 6, e27810 (2017).

52 G. Buhrman, G. Wink, and C. Mattos, Structure 15, 1618 (2007).

53 S. D. Ramalho, C. K. Wang, G. J. King, K. A. Byriel, Y.-H. Huang, V. S. Bolzani, and D. J. Craik, Journal of natural products 81, 2436 (2018).

54 T. C. Terwilliger, G. Bunkóczi, L.-W. Hung, P. H. Zwart, J. L. Smith, D. L. Akey, and P. D. Adams, Biological Crystallography 72, 359 (2016).

55 L. Di Costanzo, F. Forneris, S. Geremia, and L. Randaccio, Biological Crystallography 59, 1435 (2003).

56 C. D. Hutchison, J. M. Baxter, A. Fitzpatrick, G. Dorlhiac, A. Fadini, S. Perrett, K. Maghlaoui, S. B. Lefevre, V. Cordon-Preciado, J. L. Ferreira, et al., Nature chemistry 15, 1607 (2023).

57 A. S. Disa, T. F. Nova, and A. Cavalleri, Nature Physics 17, 1087 (2021).

58 https://github.com/Hekstra-Lab/regroup, https://github.com/Hekstra-Lab/regroup.

